# Kainate receptors coordinate primitive hematopoiesis and hemogenic niche organization to promote hematopoietic stem cell development

**DOI:** 10.64898/2026.09.21.753284

**Authors:** Margarita Parada-Kusz, Samuel J. Wattrus, Jacquelyn Jacobs, Amber D. Ide, Jack Y. Shigeta, Vincenzo di Natale, Mohamed T. Elaswad, Zhen Fu, Emily R. Goering, Stephanie Blackwood, Travis Walton, Anne E. Clatworthy, Stephanie Grainger, Deborah T. Hung

## Abstract

The adult hematopoietic system is established during embryogenesis through the generation of hematopoietic stem and progenitor cells (HSPCs) within specialized transient niches, yet the signaling mechanisms that coordinate this process remain poorly understood. Here, we identify a developmental role for kainate receptors (KARs), glutamate-gated ion channels classically associated with excitatory neurotransmission, in coordinating primitive hematopoiesis and hemogenic niche formation in zebrafish. Loss of the KAR subunits encoded by *grik1b* or *grik5* depletes primitive blood cells and disrupts hemogenic niche organization. These defects are accompanied by endothelial oxidative stress and impaired Notch signaling, which together reduce HSPC emergence and limit the clonal diversity of the adult hematopoietic stem cell pool. Shifting primitive hematopoietic output away from erythropoiesis and toward myelopoiesis restored macrophage niche colonization, endothelial Notch activity, and HSPC formation in KAR-deficient embryos. Moreover, hemogenic cell-specific Notch reactivation rescued HSPC production, demonstrating that KAR-dependent primitive hematopoiesis regulates this process through endothelial Notch signaling. Lastly, conserved KAR expression in endothelial and hematopoietic populations across zebrafish, mouse, and human suggests that these receptors may regulate blood development across multiple cellular components. Together, these findings reveal that glutamate receptor signaling extends beyond its well-established functions in neuronal transmission to coordinate the developmental processes that establish the blood system.

---

The adult hematopoietic system is established during embryogenesis through the generation of hematopoietic stem and progenitor cells (HSPCs) within a specialized embryonic microenvironment known as the hemogenic niche. In vertebrates, developmental hematopoiesis begins with a primitive wave that generates early erythroid and myeloid cells. Primitive macrophages and neutrophils derived from this wave support HSPC emergence within the developing hemogenic niche through inflammatory signaling and tissue remodeling^1–3^. Within this niche, hemogenic endothelial cells undergo an endothelial-to-hematopoietic transition to generate HSPCs that sustain lifelong hematopoiesis. Primitive macrophage–HSPC interactions further shape the clonal diversity of adult hematopoiesis by monitoring cellular stress in nascent HSPCs^4^. Yet despite the lasting consequences of these early developmental interactions for adult hematopoiesis, the signals that enable primitive hematopoietic cells to organize the hemogenic niche and support HSPC emergence remain incompletely defined^5–7^.

Kainate receptors (KARs) are evolutionarily conserved glutamate-gated ion channels that regulate synaptic transmission and neuronal circuit maturation in vertebrates. Despite their well-established neuronal functions, understanding of KAR function outside the nervous system is limited. We recently identified a role for KARs in innate immunity beyond their known canonical functions in neurotransmission, by demonstrating that chemical inhibition of KARs enhances macrophage-mediated control of bacterial infection in zebrafish embryos^8,9^. Because host defense at this stage of development is mediated by the same primitive macrophages and neutrophils that support HSPC emergence, we wondered whether KARs might regulate not only antimicrobial function but also the developmental activities of these primitive immune cells during hematopoietic development.

Here, using zebrafish (*Danio rerio*) to model hematopoietic development, we identify KAR signaling as a mechanism that coordinates primitive erythromyeloid development and hemogenic niche organization to support HSPC emergence. These findings reveal a previously unrecognized developmental role for glutamate receptor signaling in coordinating blood system formation and establishing the stem cell pool that sustains lifelong hematopoiesis.

## Results

### KARs are required for primitive immune cell development

KARs are a class of ionotropic glutamate receptors that assemble as tetramers from five subunits (GluK1–5) encoded by five distinct genes (*GRIK1*–5) in mammals. KAR subunits fall into two functionally distinct groups: GluK1–3 are principal subunits that can form functional homomeric or heteromeric receptors, whereas the higher-affinity GluK4 and GluK5 subunits require co-assembly with GluK1–3 to form functional channels^10–13^. The zebrafish genome contains seven KAR genes, including two orthologues of *GRIK1* (*grik1a* and *grik1b*), *grik2–5,* and an atypical gene, *grik-l* (*grik*-like), which lacks the amino terminal domain and has no mammalian ortholog (Supplementary Fig. 1a). Consistent with these two major KAR subfamilies, phylogenetic analysis placed the subunits encoded by *grik1a* and *grik1b* within the GluK1–3 clade and those encoded by *grik4* and *grik5* within the GluK4–5 clade (Supplementary Fig. 1b).

Because our previous work implicated GluK1-containing KARs in macrophage-dependent innate immune defense^8,9^ and pathways that regulate immune cell function can also influence lineage formation at this stage of development^14,15^, we asked whether KAR signaling could also regulate immune cell development in zebrafish embryos. We focused on receptors containing GluK1a and GluK1b, as well as GluK5, the more broadly expressed high-affinity KAR subunit that co-assembles with GluK1 *in vivo*^13,16–18^. We quantified immune cell numbers after CRISPR/Cas9-mediated gene disruption or morpholino-mediated translation inhibition of *grik1* and *grik5* by live imaging of *mpeg1:mCherry;mpx:GFP* larvae, in which macrophages and neutrophils express mCherry and GFP, respectively (Supplementary Fig. 1c–i). We found that knockdown of *grik1b* or *grik5* by CRISPR or morpholino significantly reduced the numbers of both cell types, mirroring the phenotype observed after *pu.1* knockdown, a master regulator of myelopoiesis^19^ (Fig. 1a–e and Supplementary Fig. 2a–g). CRISPR-mediated *grik1a* knockdown resulted in a milder reduction in immune cell numbers (Supplementary Fig. 2h, i). We therefore focused subsequent mechanistic studies on *grik1b* and *grik5*.

**Figure 1.**
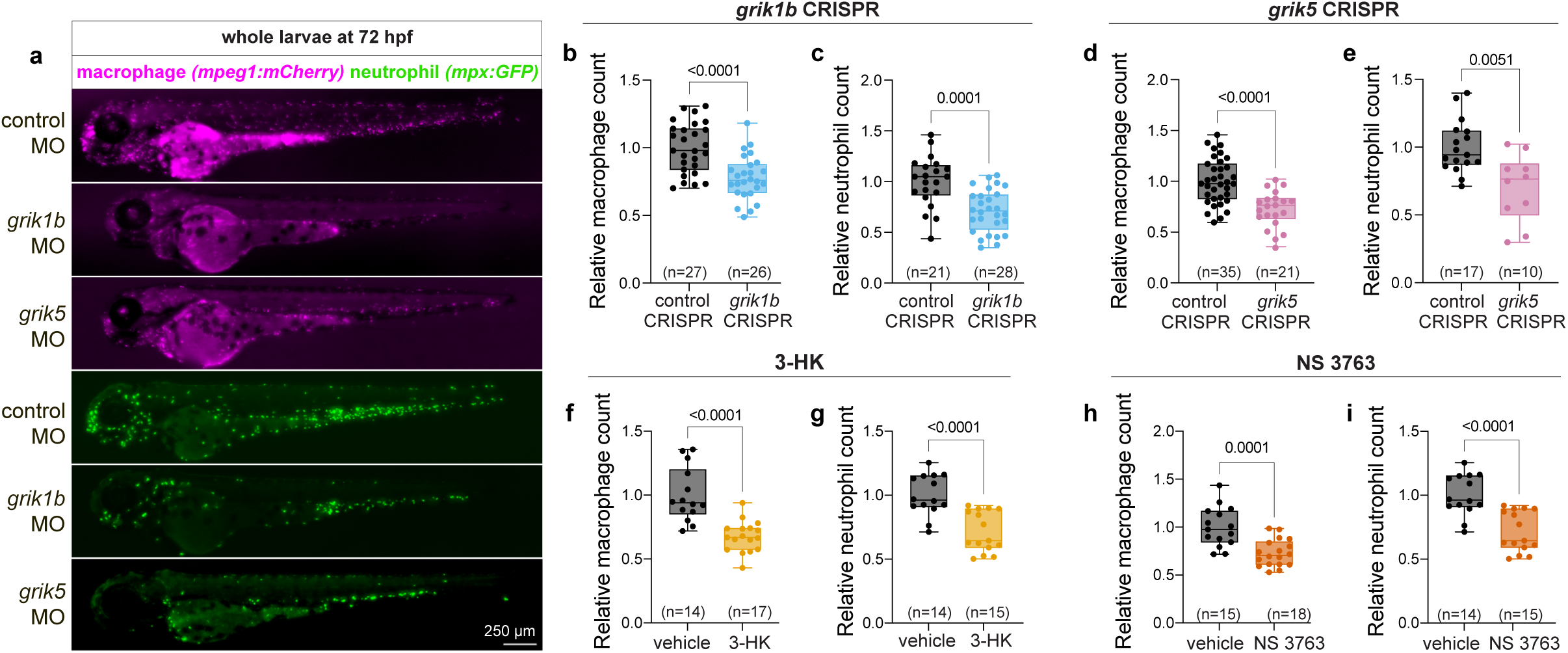
KAR knockdown or inhibition impairs primitive myeloid development. **a** Representative epifluorescence whole-larval Z-stack projections at 72 hpf of control, *grik1b*, and *grik5* morphant larvae. Macrophages are visualized using *mpeg1:mCherry* and pseudocolored magenta; neutrophils are visualized using *mpx:GFP*. **b–e** Knockdown of *grik1b* or *grik5* significantly decreases macrophage and neutrophil numbers in 72 hpf larvae. Quantification of macrophage and neutrophil cell numbers from z-stack projections of *grik1b* larvae (**b,c**) and *grik5* larvae (**d,e**). **f–i** Treatment with KAR inhibitors from 5-72 hpf also inhibits macrophage and neutrophil numbers at 72 hpf relative to controls. Larvae were treated with 3-HK (75 μM; **f,g**) or NS 3763 (75 μM; **h,i**) and cell numbers were quantified as in **b–e**. Data are shown as box-and-whisker plots with individual values; center line, median; whiskers, min to max. Statistics: two-group comparisons were performed using Welch’s unpaired t test for panels **b–f**, Student’s unpaired t test for panel **h**, and the Mann–Whitney test for panels **g** and **i**.

We confirmed these results by treating *mpeg1:mCherry;mpx:GFP* embryos early at 5 hpf with small molecule KAR antagonists 3-hydroxykynurenine (3-HK), a GluK1-containing KAR antagonist^8^, and NS 3763, a GluK1-selective noncompetitive antagonist^20^, until 72 hpf (Supplementary Fig. 1c). Quantification of immune cell numbers by live epifluorescence imaging revealed that early chemical inhibition of KARs also decreased immune cell counts when compared to vehicle treated control larvae (Fig. 1f–i). The genetic and chemical experiments together demonstrate that KARs are required for the normal development of primitive innate immune cells early in embryogenesis.

One of the earliest steps in primitive myeloid cell development is the emergence of *pu.1*-expressing progenitors, which arise predominantly in the anterior blood island, with additional progenitors arising in the posterior blood island before differentiating into macrophages and neutrophils^21^. To determine whether KARs are required for myeloid progenitor formation, we quantified *pu.1⁺* cells in the anterior and posterior blood islands by *in situ* hybridization chain reaction at 24 hpf. No significant differences were observed in either region following *grik1b* or *grik5* knockdown (Supplementary Fig. 2j–l), suggesting that early myeloid progenitor generation is largely preserved. Thus, KARs are dispensable for early progenitor formation but are required for the subsequent maintenance or expansion of primitive immune cell populations.

### KARs support primitive erythropoiesis and hemogenic niche organization

Having established that KARs regulate primitive innate immune cell development, we next asked whether they also affect primitive erythropoiesis by examining erythrocytes in *gata1a:DsRed* reporter larvae. Flow cytometry of dissociated larvae at 72 hpf revealed a significant reduction in erythrocyte abundance following *grik1b* and *grik5* knockdown, paralleling the depletion of primitive myeloid cells observed at this stage (Fig. 2a, Supplementary Fig. 3a, and Fig. 1a–e). However, in contrast to the preserved myeloid progenitor population at 24 hpf, the percentage of erythrocytes was already significantly reduced following *grik5* knockdown at this early timepoint (Fig. 2b), indicating that *grik5* loss affects erythroid development at an earlier stage. Primitive erythrocytes arise in the posterior intermediate cell mass and, upon entering the circulation, normally distribute from the axial vessels to anterior vascular territories^22^. At 24–26 hpf, using confocal imaging, we found that in contrast to control larvae which displayed this expected pattern, with erythrocytes distributed throughout the embryonic vasculature including the anterior circulation, the erythrocytes in larvae in which *grik1b* or *grik5* had been knocked down were largely confined to posterior axial vessels and markedly depleted from the anterior embryo (Fig. 2c).

**Figure 2.**
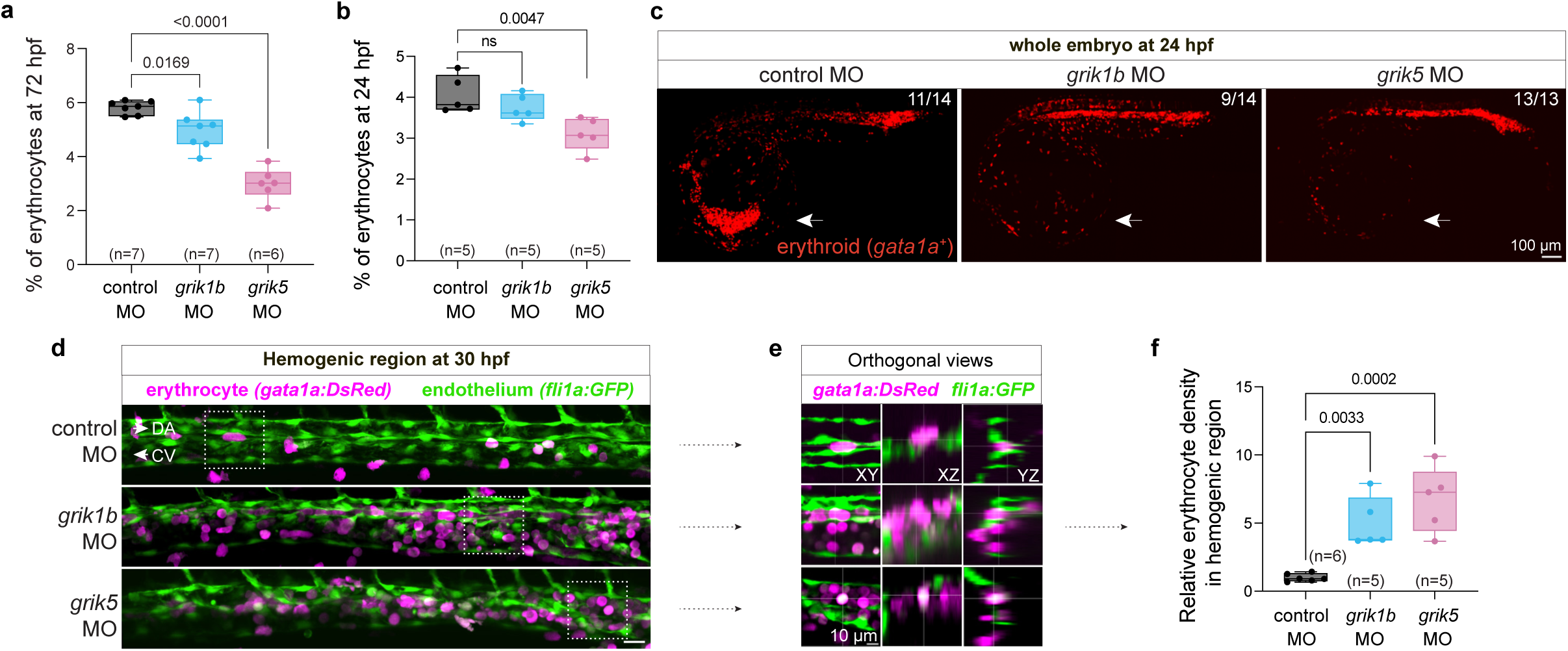
KAR knockdown impairs primitive erythroid development and disrupts erythrocyte distribution in the hemogenic region. **a,b** Quantification of the percentage of DsRed⁺ erythrocytes in control, *grik1b*, and *grik5* morphants determined by flow cytometry in *gata1a* larvae at 72 hpf (**a**) and embryos at 24 hpf (**b**). Knockdown of *grik1b* or *grik5* significantly decreases the percentage of erythrocytes at 72 hpf, while *grik5* knockdown decreases this percentage even at 24 hpf. Each dot represents one biological replicate composed of five pooled embryos or larvae. **c** Representative whole-embryo confocal z-stack projections of *gata1a* embryos showing erythroid cells in control, *grik1b*, and *grik5* morphants at 24 hpf. Erythrocytes accumulate posteriorly and are depleted from the anterior region (arrows) following *grik1b* or *grik5* knockdown compared with control morphants. Fractions indicate the number of embryos showing the depicted phenotype over the total number examined. **d** Representative confocal z-stack projections of the hemogenic region in *fli1a:GFP;gata1a:DsRed* control, *grik1b*, and *grik5* morphant embryos at 30 hpf. Knockdown of *grik1b* or *grik5* results in abnormal morphology of the dorsal aorta (DA) and caudal vein (CV), accompanied by increased erythrocyte abundance throughout the axial vasculature and adjacent subaortic space. Erythrocytes are pseudocolored magenta; endothelium is shown in green. Arrows indicate the direction of blood flow; dashed boxes indicate regions shown at higher magnification in **e**. Scale bar, 20 μm. **e** Higher-magnification confocal optical sections from the boxed regions in **d**, with orthogonal views showing erythrocyte localization relative to the axial vessels. Scale bar, 10 μm. **f** Quantification of relative erythrocyte density in the hemogenic region at 30 hpf from confocal images as shown in **d**. Data are shown as box-and-whisker plots with individual values; center line, median; whiskers, min to max. Statistics: panels **a**, **b**, and **f**, one-way ANOVA with Dunnett’s multiple-comparisons test.

Because these abnormalities could reflect altered organization of the developing vascular microenvironment rather than a primary erythroid defect, we next examined endothelial development and vascular morphology. Live imaging and flow cytometry of endothelial reporter *flk1:mCherry* embryos revealed no gross abnormalities in endothelial cell abundance or overall vascular architecture (Supplementary Fig. 3b–d), suggesting that early endothelial development is largely preserved. However, high-resolution confocal imaging of dual vascular and erythrocyte reporter *fli1a:GFP;gata1a:DsRed* embryos at 30 hpf revealed abnormal morphology of the dorsal aorta and caudal vein, accompanied by aberrant erythrocyte localization throughout the axial vasculature and adjacent subaortic space in KAR-deficient embryos (Fig. 2d–f). Because these abnormalities localize to the developing hemogenic niche, these results suggested that KAR deficiency might disrupt organization of the embryonic vascular microenvironment in which HSPCs emerge, rather than causing a primary defect in erythropoiesis alone. Although impaired cardiac function and circulation could also produce abnormal erythrocyte localization during early embryogenesis^23,24^, heart rate was unchanged in both *grik1b* and *grik5* knockdown embryos (Supplementary Fig. 3e) and, while circulation was slightly reduced in *grik5* embryos, it was unaffected in *grik1b* knockdown embryos (Supplementary Fig. 3f), indicating that cardiovascular dysfunction does not account for the vascular abnormalities shared by both KAR-deficient embryos. Collectively, these findings indicate that KARs are required not only for normal primitive hematopoietic development but also for proper organization of the embryonic hemogenic niche that supports subsequent HSPC emergence.

### KARs are required for HSPC production at the hemogenic niche

In zebrafish, HSPCs arise from specialized endothelial cells within the hemogenic niche through an endothelial-to-hematopoietic transition beginning around 26 hpf and continuing through approximately 48 hpf^25–29^. Primitive macrophages are functionally active by 24 hpf and, together with later-emerging neutrophils, support this process through inflammatory signaling and tissue remodeling^1–3,30^. After entering circulation, HSPCs seed the caudal hematopoietic tissue, a vascularized fetal niche analogous to the mammalian fetal liver^25,31^, where macrophages promote HSPC expansion and selective attrition before HSPCs colonize the adult kidney marrow and thymus to sustain adult blood production (Supplementary Fig. 4a)^4,25,31^.

Because KAR deficiency disrupted primitive hematopoiesis and altered the developing hemogenic niche, we next asked whether HSPC development was impaired. We first quantified HSPCs in the caudal hematopoietic tissue of 72 hpf larvae following KAR knockdown or chemical inhibition. Live imaging of transgenic zebrafish expressing the HSPC reporters *runx1+23:mCherry* or *runx1:eGFP* revealed that morpholino- or CRISPR-mediated knockdown of *grik1b* or *grik5* significantly reduced HSPC numbers compared to control siblings (Fig. 3a–d and Supplementary Fig. 4b). Early, continuous KAR inhibition using 3-HK or NS 3763 from 5 to 72 hpf similarly reduced HSPC numbers in the caudal hematopoietic tissue (Fig. 3e, f). These results indicate that KAR function is required for normal HSPC development.

**Figure 3.**
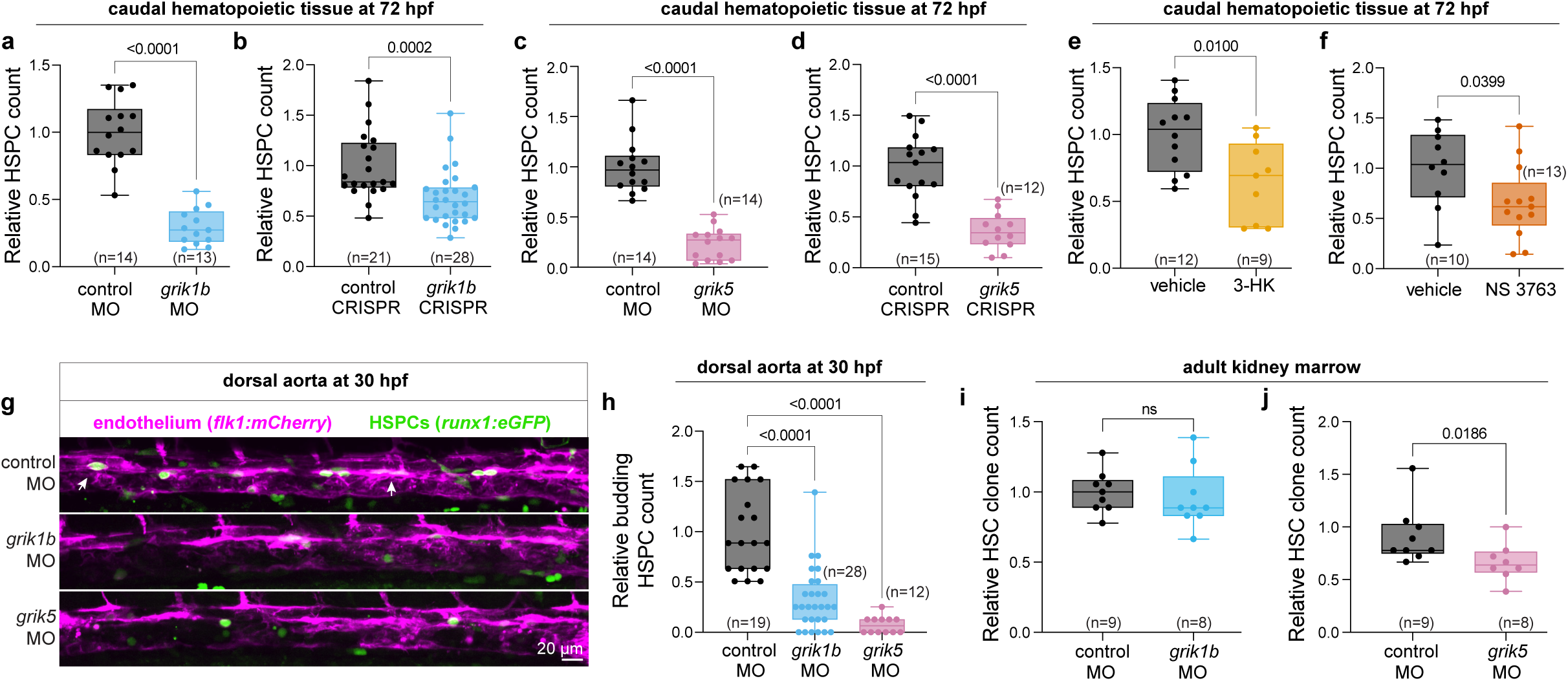
Loss of KAR function impairs HSPC development and affects adult hematopoietic clonal diversity. **a–d** Knockdown of *grik1b* or *grik5* using morpholinos (**a**, **c**) or CRISPR (**b**, **d**) significantly decreases HSPCs in 72 hpf larvae. HSPC abundance in the caudal hematopoietic tissue at 72 hpf was quantified by counting fluorescently labelled HSPCs in *runx1:eGFP* or *runx1+23:mCherry* larvae. **e**, **f** Treatment with the KAR inhibitors 3-HK (100 μM; **e**) or NS 3763 (75 μM; **f**) from 5 to 72 hpf decreases HSPC abundance in the caudal hematopoietic tissue at 72 hpf. **g**, **h** Knockdown of *grik1b* or *grik5* decreases HSPCs in the dorsal aorta at 30 hpf. **g** Representative confocal maximum-intensity z-stack projections of the dorsal aorta at 30 hpf in *flk1:mCherry;runx1:eGFP* embryos show reduced HSPC budding in *grik1b* and *grik5* morphants. Arrowheads indicate representative budding HSPCs. Endothelial cells are pseudocolored magenta. **h** Quantification of relative budding HSPC count at the dorsal aorta at 30 hpf in control, *grik1b*, and *grik5* morphants, determined from confocal z-stacks of *flk1:mCherry;runx1:eGFP* embryos and normalized to controls. **i**, **j** Transient embryonic knockdown of *grik5* (**j**), but not *grik1b* (**i**), significantly reduces HSC clonality in adult kidney marrow. Each point represents one adult fish. Data are shown as box-and-whisker plots with individual values; center line, median; whiskers, min to max. Statistics: panels **a**, **c–f**, and **i**, unpaired Student’s *t*-test; panels **b** and **j**, Mann–Whitney test; panel **h**, Kruskal–Wallis test with Dunn’s multiple-comparisons test.

The reduction in HSPCs in the caudal hematopoietic tissue at 72 hpf after loss of KAR function could reflect impaired HSPC production, survival, or proliferation. We first assessed HSPC survival and found no increase in TUNEL-positive HSPCs in the caudal hematopoietic tissue of *grik1b* or *grik5* morphants, suggesting that increased cell death is unlikely to account for the reduction in HSPC abundance (Supplementary Fig. 4c, d). We next evaluated HSPC proliferation using EdU incorporation. *grik1b* or *grik5* morphants exhibited a significant decrease in the fraction of HSPCs incorporating EdU, indicating reduced proliferation and consistent with previous studies showing that reduced primitive macrophage abundance limits HSPC expansion (Supplementary Fig. 4e, f)^4^.

We next asked whether KARs are also required for HSPC emergence from hemogenic endothelium. To test this, we used live confocal imaging to quantify HSPCs emerging from the dorsal aorta in *runx1:eGFP;flk1:mCherry* embryos, in which endothelial-to-hematopoietic transition can be visualized as budding of double-positive nascent HSPCs from the hemogenic endothelium in the ventral wall of the dorsal aorta ^26,27^. High resolution live imaging of this area revealed a marked reduction in HSPC budding following morpholino knockdown of *grik1b* or *grik5* at 30 hpf (Fig. 3g, h; Supplementary Movies 1–3). Together, these findings indicate that *grik1b* and *grik5* knockdown impair both HSPC emergence at the hemogenic niche and subsequent proliferative expansion in the caudal hematopoietic tissue.

Lastly, we asked whether the reduction in HSPC development caused by embryonic KAR deficiency has lasting consequences for the clonal composition of adult hematopoiesis. To determine whether transient disruption of KAR function alters the pool of embryonic HSC clones that persist into adulthood, we used TWISTR (tissue editing with inducible stem cell tagging via recombination)^32^ to combine morpholino-mediated knockdown of *grik1b* or *grik5* with inducible Zebrabow color labeling of hematopoietic stem cells (HSCs). In *Zebrabow-M;drl:CreERT2* embryos, individual HSC clones were permanently labeled with distinct heritable fluorescent hues beginning at 24 hpf, allowing for later analysis of hematopoietic clonality through adult marrow dissection (Supplementary Fig. 4g)^33,34^. Analysis of adult marrow granulocytes revealed significantly reduced hematopoietic clonal diversity following *grik5* knockdown, with a similar trend following *grik1b* knockdown (Fig. 3i, j and Supplementary Fig. 4h). These findings indicate that transient disruption of KAR function during embryogenesis can reduce the number of embryonic HSC clones contributing to adult hematopoiesis.

### Redirecting primitive hematopoiesis rescues impaired HSPC formation in KAR-deficient embryos

Because primitive hematopoietic defects preceded impaired HSPC production and expansion, we next tested whether correcting hemogenic niche composition could rescue HSPC emergence. To simultaneously reduce erythrocyte accumulation and increase primitive myeloid cells in *grik1b*- and *grik5*-morphant embryos, we co-injected a morpholino targeting *gata1a*, which suppresses erythroid specification and redirects primitive hematopoietic output toward myelopoiesis ^19,35^. Confocal imaging of *mpeg1:mCherry;mpx:GFP* larvae revealed that *gata1a* co-knockdown restored macrophage and neutrophil abundance in both KAR morphants (Supplementary Fig. 5a–d).

We next examined whether *gata1a*-mediated restoration of immune cell numbers was accompanied by increased macrophage abundance near the hemogenic region, where macrophages support HSPC mobilization and hemogenic niche remodeling^3,36^. Quantification within the axial vasculature spanning the dorsal aorta and caudal vein of *mpeg1:mCherry;fli1a:GFP* embryos at 48 hpf revealed reduced macrophage abundance in KAR morphants, which was restored by co-knockdown of *gata1a* with *grik1b* or *grik5* (Fig. 4a–c). We then asked whether this restoration was sufficient to rescue HSPC emergence. Indeed, *gata1a* co-knockdown normalized both the number of budding *runx1*-positive HSPCs emerging from the hemogenic endothelium (Fig. 4d–f) and the downstream accumulation of *runx1:eGFP*⁺ HSPCs in the caudal hematopoietic tissue of KAR-deficient larvae (Supplementary Fig. 5e, f). Collectively, these data indicate that impaired HSPC emergence in KAR-deficient embryos arises from defective primitive hematopoietic development and that shifting primitive hematopoietic output away from erythropoiesis and toward myelopoiesis is sufficient to replenish the primitive immune cell component of the hemogenic niche and rescue HSPC emergence.

**Figure 4.**
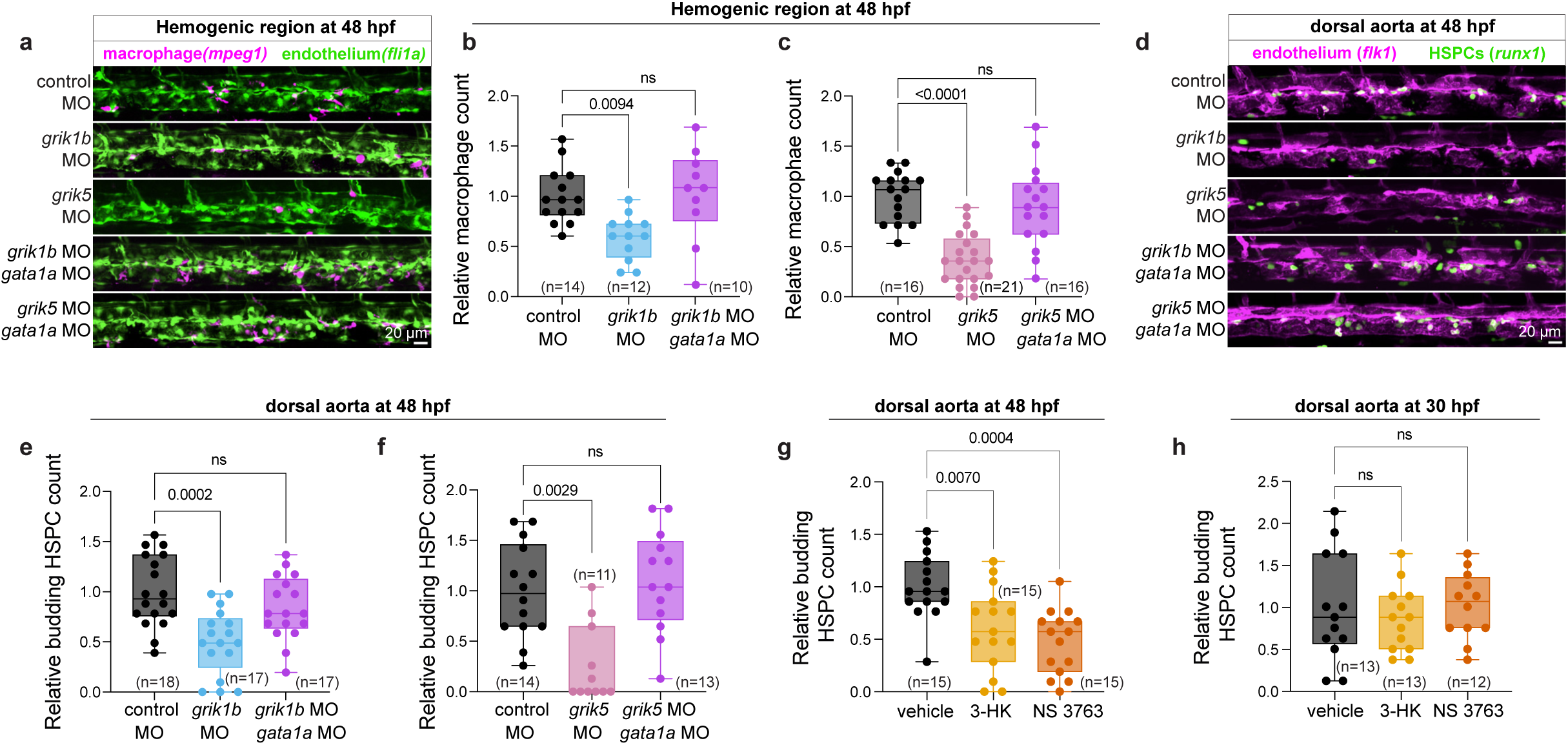
Redirecting primitive hematopoietic output toward myelopoiesis restores macrophage colonization and HSPC emergence in KAR-deficient embryos. **a** Representative confocal z-stack projections of the hemogenic region in *mpeg1:mCherry;fli1a:GFP* embryos in control, *grik1b*, *grik5*, *grik1b* + *gata1a*, and *grik5* + *gata1a* morphants; macrophages are pseudocolored magenta. **b,c** Quantification of macrophage numbers from confocal projections shows that *gata1a* co-knockdown restores macrophage numbers in the hemogenic niche at 48 hpf relative to the corresponding *grik1b* or *grik5* single morphants. **d** Representative confocal z-stack projections of the dorsal aorta at 48 hpf in *flk1:mCherry;runx1:eGFP* embryos in control, *grik1b*, *grik5*, *grik1b* + *gata1a*, and *grik5*+ *gata1a* morphants; endothelial cells are pseudocolored magenta. **e,f** Quantification of budding HSPCs shows that *gata1a* co-knockdown restores budding HSPC numbers in the dorsal aorta at 48 hpf relative to the corresponding *grik1b* or *grik5* single morphants. **g,h** KAR inhibition with 3-HK (100 μM) or NS 3763 (75 μM) from 5 to 24 hpf reduces budding HSPC counts at 48 hpf (**g**), whereas inhibition from 24 to 30 hpf does not significantly alter budding HSPC counts at 30 hpf (**h**), relative to vehicle-treated controls. Data are shown as box-and-whisker plots with individual values; center line, median; whiskers, min to max. Statistics: panels **b**, **c**, **e**, **g**, and **h**, one-way ANOVA with Šídák’s multiple-comparisons test; panel **f**, Kruskal–Wallis test with Dunn’s multiple-comparisons test.

The *gata1a* rescue experiments suggested that KARs promote HSPC emergence indirectly by regulating primitive hematopoietic development. If so, KAR activity should be required during the early developmental window encompassing primitive hematopoiesis and hemogenic niche formation, rather than later during HSPC emergence itself. To test this, we chemically inhibited KAR activity during discrete developmental windows using 3-HK or NS 3763. Transient inhibition of KAR activity with 3-HK or NS 3763 during the first 24 hpf, when primitive blood cells and the axial vasculature are established, was sufficient to reduce the number of budding HSPCs at 48 hpf (Fig. 4g). In contrast, inhibition initiated at 24 hpf, after primitive hematopoiesis was established, had no impact on HSPC emergence at 30 hpf (Fig. 4h). These results indicate that the critical window for KAR activity precedes HSPC emergence and coincides with primitive hematopoietic development and hemogenic niche formation, further supporting a model in which KARs promote HSPC emergence indirectly by early establishment of the primitive hematopoietic environment required for hemogenic niche function.

### KARs promote HSPC emergence through endothelial Notch signaling

Having established that KAR activity is required for primitive hematopoiesis and normal hemogenic niche development, we next asked what molecular alterations arise in the endothelium of KAR-deficient embryos before HSPC emergence. We conducted bulk transcriptional profiling of endothelial cells isolated by flow cytometry from single-cell suspensions of 24 hpf *flk1:mCherry* and *flk1:GFP* embryos following control, *grik1b* or *grik5* morpholino knockdown (Supplementary Fig. 6a). Differential expression analysis revealed convergent transcriptional alterations in both knockdowns relative to controls, with *grik5* morphants exhibiting more pronounced effects, consistent with our functional data. Both *grik1b* and *grik5* knockdown induced transcriptional programs associated with FoxO and apoptotic signaling, p53-associated cellular stress and cell-cycle arrest, and suppression of DNA replication and ribosome biogenesis (Fig. 5a, b and Supplementary Fig. 6b–f). In addition, *grik5* knockdown altered transcriptional programs linked to blood circulation, consistent with the reduced number of circulating blood cells observed specifically in *grik5* but not *grik1b* morphants (Supplementary Fig. 6g and Supplementary Fig. 3f). Flow cytometric analysis further showed increased ROS-associated fluorescence in endothelial cells from *grik1b* and *grik5* morphants, consistent with elevated intracellular oxidative stress (Fig. 5c and Supplementary Fig. 6j). Together, these results show that KAR deficiency drives the endothelium into a stress-associated cellular state before HSPC emergence.

**Figure 5.**
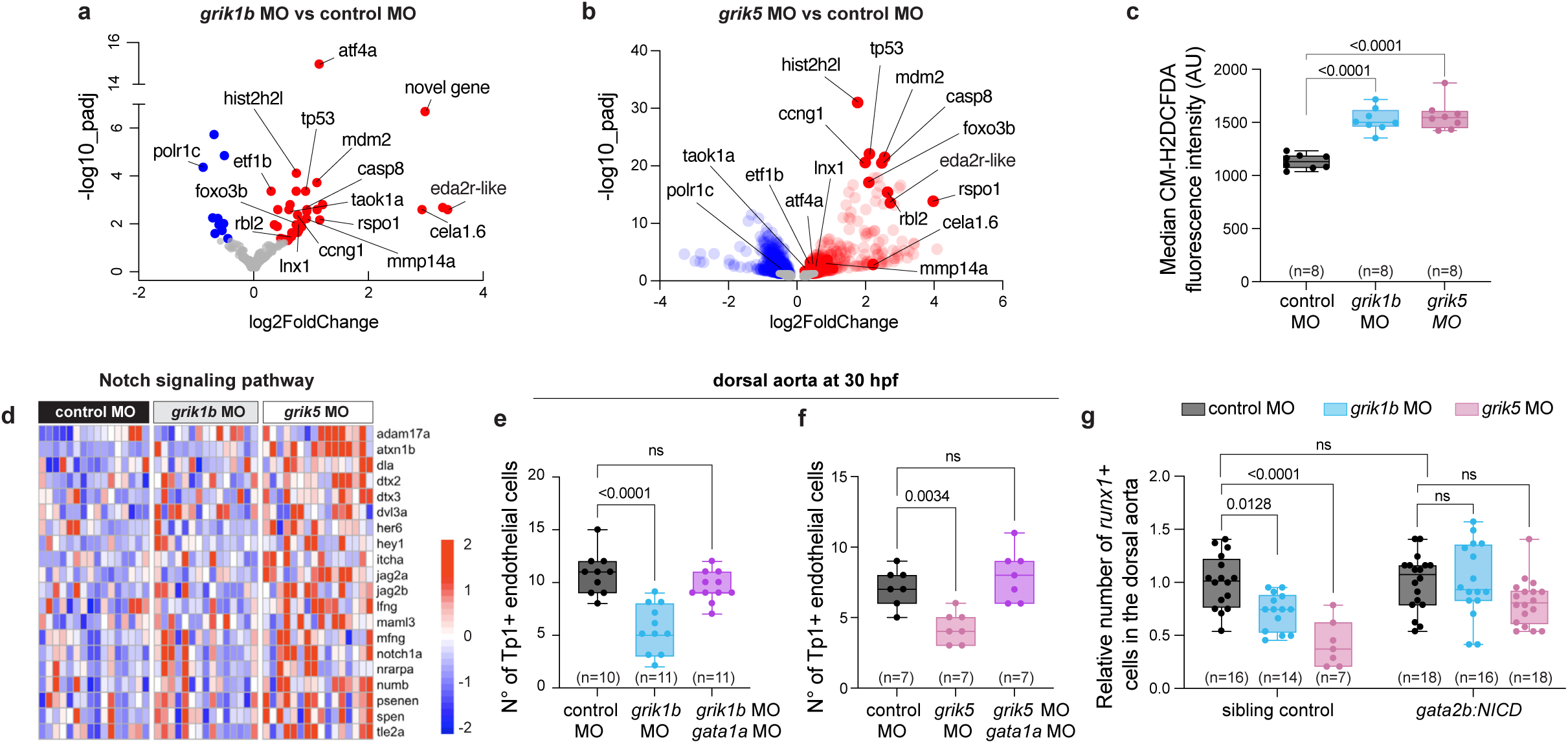
KAR deficiency induces endothelial stress and impairs Notch signaling required for HSPC emergence. **a,b** Bulk transcriptional profiling of sorted endothelial cells at 24 hpf shows differential expression of genes associated with FoxO and apoptotic signaling, p53-associated cellular stress and cell-cycle arrest, and suppression of DNA replication and ribosome biogenesis in *grik1b* (**a**) and *grik5* (**b**) knockdown embryos compared with controls. Labeled genes include shared genes changed in the same direction in both knockdowns. **c** Endothelial cells from dissociated *grik1b* and *grik5* knockdown *flk1:mCherry* embryos display increased ROS, determined by CM-H2DCFDA fluorescence and flow cytometry at 24 hpf. Each point represents one biological replicate composed of five pooled embryos. **d** Notch-related gene sets are enriched in endothelial cells in *grik1b* and *grik5* morphants compared with controls at 24 hpf. **e,f** *grik1b* (**e**) and *grik5* (**f**) morphants show a significant reduction in the number of *Tp1:mCherry*^+^ endothelial cells in the ventral wall of the dorsal aorta at 30 hpf, with higher counts observed following *gata1a* co-knockdown. **g** Higher numbers of *runx1*⁺ cells are observed along the dorsal aorta in *grik1b* and *grik5* morphants at 30 hpf following constitutive activation of Notch in the hemogenic endothelium. *runx1*⁺ cells were quantified by whole-mount in situ hybridization in control and KAR-deficient siblings with or without the *gata2b:KalTA4;UAS:NICD* transgene. Data are shown as box-and-whisker plots with individual values; center line, median; whiskers, min to max. Statistics: panels **c**, **e**, and **f**, one-way ANOVA with Dunnett’s multiple-comparisons test; panel **g**, two-way ANOVA with Šídák’s multiple-comparisons test for selected pairwise comparisons.

We also found that Notch signaling, which is essential for arterial development and subsequent HSPC emergence^28,37–39^, is dysregulated in KAR deficiency. Notch-related gene sets were enriched in both *grik1b*- and *grik5*-deficient endothelial cells, with *grik5* knockdown additionally showing enrichment of vascular development signatures (Fig. 5d and Supplementary Fig. 6c, h, i), raising the possibility that Notch-dependent hemogenic endothelial programs are altered during HSPC formation. We therefore asked whether Notch activity was changed at 30 hpf, when it is required for HSPC emergence^40^. Using *fli1a:GFP;Tp1:mCherry* embryos in which canonical Notch signaling drives *Tp1:mCherry* reporter expression^41^, we found that both *grik1b* and *grik5* morphants showed significantly reduced Tp1 reporter activity along the ventral wall of the dorsal aorta (Fig. 5e, f and Supplementary Fig. 6k), indicating diminished endothelial Notch signaling. Notably, *gata1a* co-knockdown restored Tp1 reporter activity in both backgrounds (Fig. 5e, f and Supplementary Fig. 6k). Thus, redirecting primitive hematopoietic output toward myelopoiesis restored endothelial Notch signaling and rescued HSPC emergence (Fig. 4d–f). Given that primitive neutrophils and macrophages support this process through TNFα/NF-κB–Jagged1–Notch–Runx1 signaling and extracellular matrix remodeling, respectively^1,3,36^, these results suggest that KARs promote HSPC emergence by regulating primitive hematopoietic development upstream of endothelial Notch signaling.

We tested this hypothesis by asking whether restoring Notch activity alone was sufficient to rescue HSPC emergence in KAR-deficient embryos. We used the *gata2b:KalTA4;UAS:NICD*binary transgenic system to express the constitutively active Notch1a intracellular domain (NICD) specifically in *gata2b*+ hemogenic endothelium^29^. In this system, *grik5* and *grik1b* morphant siblings lacking the NICD transgenic combination showed significantly reduced *runx1* expression along the ventral wall of the dorsal aorta, whereas KAR-deficient embryos carrying both transgenes restored *runx1* expression to control sibling levels (Fig. 5g and Supplementary Fig. 6l). These data demonstrate that constitutive activation of Notch signaling within the hemogenic endothelium is sufficient to rescue HSPC emergence.

Collectively, these findings show that KAR deficiency induces endothelial stress and impairs endothelial Notch signaling. Restoration of endothelial Notch signaling is sufficient to rescue HSPC emergence, placing endothelial Notch signaling downstream of KAR signaling in the establishment of definitive hematopoiesis.

### KAR subunits are expressed in diverse hematopoietic populations across vertebrates

Our findings identified a developmental mechanism by which KARs coordinate primitive hematopoietic development, hemogenic niche formation, and HSPC development in zebrafish. Given the well-established roles of KARs in the nervous system, we next asked where KAR signaling might act during hematopoietic development, whether in neurons, hematopoietic cells, supporting niche populations, or across multiple embryonic tissues. We therefore examined *grik1b* and *grik5* expression in zebrafish embryos and assessed whether these patterns were conserved in mouse and human single-cell transcriptomic datasets.

We first examined KAR gene expression in zebrafish embryos at 24 hpf using *in situ* hybridization chain reaction. *grik5* was broadly detected throughout the developing brain and spinal cord, whereas *grik1b* signal was restricted to a discrete region of the anterior brain and scattered yolk-associated cells (Fig. 6a, b), consistent with the known enrichment of KAR expression in neuronal tissues. *grik1b* also overlapped with a subset of *pu.1+/fli1a+* cells in the head region (Fig. 6b, arrows), consistent with our previous transcriptomic data showing *grik1b* expression in primitive macrophages^8^. Although *grik5* was not readily detected in myeloid or endothelial cells by *in situ* hybridization, both endothelial RNA-seq and an independent zebrafish single-cell atlas^42^ identified *grik5* expression in vascular, erythroid, and immune cell populations (Supplementary Fig. 7a–d). By contrast, *grik1b* showed a more restricted expression pattern across the same populations (Supplementary Fig. 7a–d).

**Figure 6.**
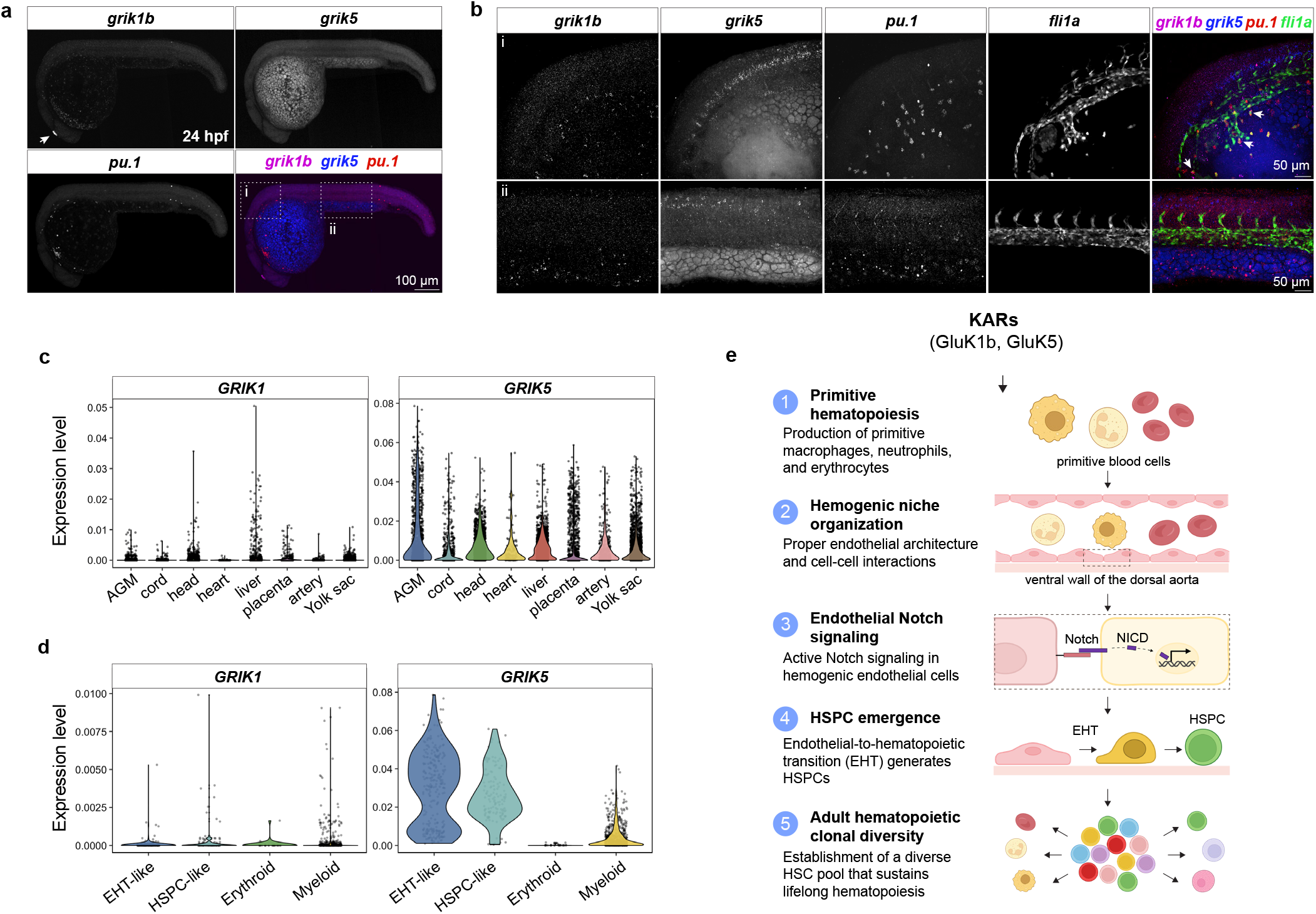
KAR subunit expression is conserved in neural and hematopoietic tissues across vertebrates. **a,b** Whole-mount hybridization chain reaction in zebrafish embryos at 24 hpf shows *grik1b* expression in a discrete region of the anterior brain (**a**, arrow, pseudocolored magenta) and scattered yolk-associated cells, whereas *grik5* broadly localizes to the developing brain and spinal cord. *grik1b* signal overlaps with a subset of *pu.1*⁺/*fli1a*⁺ cells in the head region (**b**). Dashed boxes in **a** indicate the anterior/head region (i) and trunk/hemogenic region (ii) shown at higher magnification in **b**; arrows indicate representative areas of signal overlap. Endothelial cells are visualized using the *fli1a:GFP* reporter. **c,d** Violin plots showing MAGIC-imputed *GRIK1* and *GRIK5* expression across human developmental hematopoietic tissues (**c**) and in human EHT-like, HSPC-like, erythroid, and myeloid populations (**d**), based on reanalysis of the published scRNA-seq dataset from *Calvanese et al.*, 2022^44^. **e** Model summarizing the proposed role of KAR signaling in coordinating primitive hematopoiesis, hemogenic niche organization, endothelial Notch signaling, HSPC emergence, and establishment of adult hematopoietic clonal diversity. Loss of KAR function disrupts this developmental sequence, with *grik5* knockdown additionally reducing adult hematopoietic clonal diversity. NICD, Notch intracellular domain; EHT, endothelial-to-hematopoietic transition.

To investigate whether KAR expression similarly spans multiple hematopoietic and supporting cell populations in mammals, we next examined KAR expression during mouse hematopoietic development using a time-resolved single-cell transcriptomic atlas of prenatal mouse development spanning gastrulation to birth^43^. We focused on stages E8.5–E11.5, encompassing mesodermal specification, hematoendothelial progenitor formation, hemogenic endothelium development, and HSPC emergence. *Grik5* was detected across mesodermal progenitors, hematoendothelial progenitors, arterial endothelium, HSCs, primitive erythrocytes, and macrophages, with expression varying across developmental stages (Supplementary Fig. 7e). *Grik1* expression was low and sparse across the populations examined, with the strongest enrichment in early mesodermal progenitors and neuronal populations, consistent with the comparatively low and anatomically restricted *grik1b* signal observed in zebrafish (Supplementary Fig. 7e).

Lastly, to assess whether the developmental expression patterns identified in zebrafish and mouse are also present during human hematopoiesis, we interrogated a single-cell transcriptomic atlas of human hematopoietic tissues spanning the first trimester to birth^44^. *GRIK5* was detected across the major tissues analyzed, including the aorta-gonad-mesonephros, ventral artery, yolk sac, cord blood, and fetal liver, with the most consistent expression in endothelial-to-hematopoietic transition-like and HSPC-like populations within the aorta-gonad-mesonephros (Fig. 6c, d). By contrast, *GRIK1* expression was markedly lower and largely restricted to a subset of myeloid cells within the aorta-gonad-mesonephros, consistent with expression patterns observed in mouse and zebrafish (Fig. 6c, d).

In sum, these cross-species analyses revealed KAR ortholog expression across diverse endothelial and hematopoietic populations in zebrafish, mouse, and human, including cell types associated with primitive hematopoiesis, endothelial-to-hematopoietic transition, and HSPC emergence. The broad distribution of KAR expression not only in neuronal populations but also in endothelial and hematopoietic compartments across vertebrates suggests that KAR-dependent regulation of hematopoiesis may involve signaling in multiple cellular compartments rather than within a single cell type or tissue. Taken together, these findings support a possible model (Fig. 6e) in which KAR signaling coordinates interactions among multiple developmental cell populations to promote primitive hematopoiesis, hemogenic niche development, and HSPC emergence.

## Discussion

We previously identified KARs as regulators of primitive macrophage function during infection, revealing a new connection between these ion channels and innate immune biology^8,9^. Because primitive immune cells also contribute to the developmental processes that support HSPC emergence, this finding raised the possibility that KAR signaling might have a broader role in early hematopoietic development. Here, we show that the KAR subunits GluK1b and GluK5 coordinate primitive hematopoietic development and hemogenic niche formation to promote HSPC formation, thus extending the role for KARs from innate immune cell function to the development of the hematopoietic system itself. Using complementary genetic and chemical approaches in zebrafish, we found that disruption of KAR function impaired primitive erythroid and myeloid cell development, disrupted hemogenic niche organization, attenuated Notch signaling, and ultimately reduced HSPC emergence. Rescue of immune cell abundance, endothelial Notch signaling, and HSPC emergence by *gata1a* co-knockdown, together with restoration of HSPC emergence through constitutive activation of Notch signaling in the hemogenic endothelium, demonstrates that KAR-dependent primitive hematopoiesis functions upstream of endothelial Notch signaling.

Our findings suggest that disruption of the hemogenic niche is a defining feature of the KAR-deficient phenotype. Erythrocyte accumulation in the axial vasculature and reduced macrophage colonization indicate that KAR deficiency disrupts the embryonic niche required for HSPC emergence. The mechanism of erythrocyte accumulation remains unclear but may involve impaired vascular integrity or cell migration^45^, contributing to endothelial stress in KAR-deficient embryos. Importantly, redirecting primitive hematopoietic output away from erythropoiesis and toward myelopoiesis through *gata1a* co-knockdown restored immune cell abundance and rescued HSPC emergence, identifying dysregulated primitive hematopoiesis as a major contributor to these defects. Together, these findings support a model in which disruption of primitive hematopoiesis impairs hemogenic niche formation, leading to impaired Notch signaling and HSPC emergence (Fig. 6e).

Importantly, the consequences of embryonic KAR deficiency extend beyond embryonic hematopoiesis. Transient embryonic *grik5* knockdown reduced the clonal diversity of hematopoietic stem cells contributing to the adult hematopoietic system, indicating that KARs have a long-lasting impact on HSC establishment and clonal diversity. Together with emerging evidence that variation in HSC establishment before birth can shape clonal hematopoiesis later in life^4,46^, these findings suggest that developmental KAR dysfunction may durably influence the adult hematopoietic clonal architecture, a feature altered during aging and hematological malignancy^47,48^.

In neural tissues, KARs primarily function as glutamate-gated cation channels, although they can also engage metabotropic, G protein-coupled signaling pathways^49–51^. The ability of NS 3763 and 3-HK, which inhibit KAR ion channel activity, to phenocopy the primitive hematopoietic and HSPC emergence defects caused by genetic KAR disruption demonstrates that ion channel activity is a functionally relevant component of KAR-dependent hematopoietic development. This finding is conceptually significant because it identifies ion channel activity as an additional regulatory layer in embryonic hematopoiesis, complementing the transcriptional and extracellular signaling pathways that dominate current models of hematopoietic development. While the sources of glutamate during hematopoietic development remain unknown, the established roles of KARs in neural signaling, together with accumulating evidence that the nervous system contributes to hematopoietic development and stem cell regulation^52–56^, raise the possibility that KAR-dependent regulation of hematopoiesis may be driven by glutamatergic signaling originating from local hematopoietic populations, neighboring developmental compartments, neural tissues, or multiple tissues acting together in a coordinated fashion.

Notably, the conserved expression of KAR orthologs in neuronal, endothelial, and hematopoietic populations across vertebrates suggests that KAR-dependent regulation of hematopoiesis may arise through signaling across multiple developmental cell populations rather than a single cell or tissue type. Although this expands the developmental context in which KARs are known to function, it is consistent with accumulating evidence that ionotropic glutamate receptors regulate mature blood and immune cell biology. Recent studies have described AMPA and NMDA receptors as regulators of megakaryocytic, erythroid, and lymphoid lineages^57–63^, while KAR signaling, through our work and that of others, has been implicated in macrophages, mast cells, B lymphocytes, and endothelial biology^8,9,64–66^. Collectively, these observations point to a broader role for glutamatergic signaling across hematopoietic, immune, and vascular systems. Our findings extend this concept into embryonic development by identifying KAR signaling as a regulator of the coordinated processes that establish the blood system, raising the possibility that glutamatergic signaling represents a previously unrecognized mechanism for organizing development across multiple cellular and tissue compartments.

The expression of KAR orthologs across zebrafish, mouse, and human hematopoietic development is also consistent with the broad conservation of developmental and molecular mechanisms across vertebrate hematopoiesis^67,68^. Notably, *GRIK5* knockdown has been reported to reduce the proliferative output of human cord blood hematopoietic progenitors^69^. Together with prior studies showing that KARs and other ionotropic glutamate receptors modulate blood and immune cell function in mouse and human models, these findings support the possibility that KAR signaling is conserved in mammalian hematopoiesis.

In summary, this work identifies KARs as newly identified regulators of developmental hematopoiesis that coordinate primitive hematopoiesis, hemogenic niche organization, and Notch-dependent HSPC emergence. These findings extend the biological role of glutamate receptors beyond their well characterized roles in neurotransmission, while providing a new framework for understanding how ion-channel signaling shapes vertebrate blood development and function.

## Methods

### Zebrafish husbandry

Wild-type AB zebrafish and the following transgenic lines were used in this study: Tg(*mpeg1:mCherry*)(gl23Tg), hereafter referred to as *mpeg1:mCherry*^70^; Tg(*mpeg1:EGFP*)^gl22Tg^, hereafter *mpeg1:EGFP*^70^; TgBAC(*mpx:GFP*)^i114^, hereafter *mpx:GFP*^71^; Tg(*runx1+23:mCherry*) and Tg(*runx1:eGFP*)^y509^, hereafter *runx1+23:mCherry* and *runx1:eGFP*^31,72^; *flk1:mCherry*, *flk1:GFP*, and Tg(*fli1a:eGFP*)^y1Tg^, hereafter *fli1a:GFP*^73–75^; Tg(*gata1a:DsRed*)^sd2Tg^, hereafter *gata1a:DsRed*^76^; Tg(*EPV.TP1-Mmu.Hbb:nlsmCherry*)^ia7^, hereafter *Tp1:mCherry*^41^; Tg(*UAS:myc-notch1a-intra*)^kca3^, hereafter *UAS:NICD*^77^; and Tg(*gata2b:KalTA4*)^sd32Tg^, hereafter *gata2b:KalTA4* ^29^; Zebrabow-M^34^ Tg(*drl:Cre-ERT2*), hereafter *drl:CreERT2*^78^. Zebrafish were maintained in accordance with institutional guidelines and approved animal protocols at Massachusetts General Hospital and Van Andel Institute, including MGH IACUC protocol #2006N000168 and VAI animal use protocols #24-08-024 and #25-04-011.

### Small molecules

3-hydroxy-D,L-kynurenine was purchased from BOC Sciences (>98%) or Biosynth Carbosynth (98%) and dissolved in 0.1 N hydrochloric acid. NS 3763 was purchased from Tocris (99%), ChemBridge (≥90%), or was synthesized in-house, and was dissolved in DMSO. Embryos were treated with 3-HK or NS 3763 at the indicated concentrations and developmental windows, with matched HCl- or DMSO-containing vehicle controls, respectively.

### Morpholino experiments

Morpholinos (Gene Tools; for sequences, see Supplementary Table 1) were suspended in sterile water, diluted to the indicated concentrations, and microinjected into 1-4-cell-stage embryos. For knockdown of *pu.1*, *gata1a* and *grik5,* previously validated gene-specific morpholinos were used: *pu.1* MO (1 nL; 2–4 ng/nL) ^19^, *gata1a* MO (1 nL; 1.69 ng/nL)^35^ and *grik5* MO (1 nL; 0.5–0.6 mM)^66^. A custom morpholino targeting *grik1b* was used for knockdown (1 nL, 1–1.5 ng/nL). A randomized control oligo, 25-N, was used as a negative control (1 nL, 1–1.5 ng/nL). For morpholinos used within a concentration range, the final working dose was optimized within the indicated range according to transgenic background sensitivity. Oligonucleotide sequences are provided in Supplementary Table 1.

### Generation of CRISPR reagents

CRISPR reagents were prepared using either *in vitro*-transcribed single-guide RNAs (sgRNAs) and Cas9 mRNA or synthetic CRISPR–Cas9 ribonucleoprotein complexes (RNPs). The target sequence, targeted exon and guide combination used for each experiment are provided in Supplementary Table 1.

For the sgRNA–Cas9 mRNA workflow, CRISPRscan (*grik1b*; https://www.crisprscan.org/)^79^ or the sgRNA prediction tool at crispr.mit.edu^80^ (*grik1a*) was used to identify candidate target sequences. Oligonucleotides containing the T7 promoter, a 20-nucleotide target-specific sequence, and 15–23 nucleotides complementary to a constant oligonucleotide encoding the reverse complement of the tracrRNA tail were annealed. Single-stranded regions were filled in using T4 DNA polymerase (NEB), and the resulting sgRNA DNA templates were purified using QIAquick PCR Purification Columns (Qiagen), as previously described^81^. sgRNAs were transcribed using the MEGAscript T7 Transcription Kit (Ambion), treated with DNase, and purified by ethanol–ammonium acetate precipitation. sgRNA concentration was measured using a NanoDrop spectrophotometer, and RNA integrity was assessed using a TapeStation system (Agilent) or a 6% TBE–urea polyacrylamide gel.

Capped Cas9 mRNA was transcribed from the pT3TS-nCas9n plasmid (a gift from Wenbiao Chen; Addgene plasmid 46757)^82^. Briefly, 2 µg of plasmid was linearized with XbaI and treated with 200 µg/mL proteinase K and 0.5% SDS for 1 h at 50 °C, followed by incubation at 95 °C for 10 min. The linearized plasmid was purified by phenol–chloroform extraction and ethanol precipitation with 5 M ammonium acetate. One microgram of purified template was transcribed using the mMESSAGE mMACHINE T3 Transcription Kit (Invitrogen) for 2 h at 37 °C. The reaction was treated with TURBO DNase for 15 min at 37 °C, and Cas9 mRNA was purified by phenol–chloroform extraction and ethanol precipitation. Cas9 mRNA concentration was measured using a NanoDrop spectrophotometer, and RNA integrity was assessed using a TapeStation system.

For the synthetic RNP workflow, two target-specific Alt-R CRISPR–Cas9 crRNAs for *grik1a* and three each for *grik1b* and *grik5*, together with tracrRNA, were obtained from Integrated DNA Technologies and prepared according to the F0 knockout method previously described^83^. Briefly, each crRNA and the tracrRNA were resuspended at 200 µM in IDT Duplex Buffer. For each target, 1 µL of crRNA, 1 µL of tracrRNA, and 1.28 µL of Duplex Buffer were incubated at 95 °C for 5 min. One microliter of the resulting 61 µM crRNA:tracrRNA duplex was combined with 1 µL of 61 µM recombinant Cas9 protein and incubated at 37 °C for 5 min. The three RNPs targeting each gene were then pooled.

### Generation of grik1a, grik1b, and grik5 F0 crispants

For experiments using in vitro-transcribed sgRNAs, approximately 50–100 pg of each sgRNA and approximately 600 pg of capped Cas9 mRNA were microinjected into the blastomere of one-cell-stage embryos. For experiments using synthetic CRISPR–Cas9 RNPs, pooled complexes were injected into one-cell-stage embryos as previously described^83,84^. Guide identities and target sites are provided in Supplementary Fig. 1 and Supplementary Table 1. Embryos injected with control guides or uninjected clutch-matched embryos served as controls.

### Assessment of CRISPR editing efficiency

At 3 days post-fertilization, individual crispant and corresponding control larvae were euthanized for genomic DNA isolation. Each larva was incubated in 100 µL of 50 mM NaOH for 20 min at 95 °C, immediately placed on ice, and neutralized with 10 µL of 1 M Tris-HCl, pH 7.5. Regions spanning each CRISPR target site were amplified using Q5 High-Fidelity DNA Polymerase (NEB) and the primer pairs listed in Supplementary Table 1. PCR products were purified using QIAquick PCR Purification Columns and subjected to paired-end amplicon sequencing by the MGH CCIB DNA Core. Editing frequency and the distribution of insertion and deletion alleles at each target site were quantified using CRISPResso2^85^.

### Assessment of transcript abundance in F0 crispants by RT-qPCR

RNA was isolated at 3 days post-fertilization from pools of eight larvae using the RNeasy Mini Kit (Qiagen) with on-column DNase treatment. Larvae were homogenized in 350 µL of RLT buffer by eight passages through a 17-gauge, 1.5-inch needle attached to a 1-mL syringe. RNA was eluted in 30 µL of nuclease-free water, and its concentration and purity were measured using a NanoDrop spectrophotometer. cDNA was synthesized from 2 µg of RNA using oligo(dT)20 primers and SuperScript IV reverse transcriptase (Life Technologies).

The relative abundance of *grik1a*, *grik1b*, and *grik5* transcripts was measured in the corresponding crispant larvae relative to the corresponding clutch-matched controls. *grik1a* was amplified using primers SB139 and SB140, *grik1b* using AC1006 and AC1007, and *grik5* using VD_33 and VD_34. Expression was normalized to *eef1a1l1* using primers AC1002 and AC1003. RT-qPCR reactions contained 1× iTaq Universal SYBR Green Supermix (Bio-Rad), 417 nM of each primer, and cDNA corresponding to 50 ng of input RNA in a final volume of 15 µL. Reactions were incubated at 95 °C for 3 min, followed by 40 cycles of 95 °C for 30 s, 54 °C for 30 s, and 72 °C for 30 s using a Bio-Rad CFX384 system. Samples were analyzed in four technical replicates, and relative expression was calculated using the ΔΔCt method.

### Embryo dissociation and fluorescence-activated cell sorting (FACS)

Single-cell suspensions were prepared from transgenic embryos at the indicated developmental stages for flow cytometry or fluorescence-activated cell sorting. For flow-cytometric analyses, embryos or larvae were collected in groups of five and dissociated enzymatically into single-cell suspensions. Endothelial samples used for RNA sequencing were prepared separately as described below. *gata1a:DsRed* embryos were dissociated for flow-cytometric analysis of erythrocytes at 24 or 72 hpf. *flk1:mCherry* and *flk1:GFP* embryos were dissociated at 24 hpf, and fluorescent endothelial cells were isolated by FACS for downstream RNA sequencing. At 24 hpf, samples were dissociated using 0.5% trypsin-EDTA (Thermo Fisher, 15400054), whereas 72-hpf samples were dissociated using 0.5% trypsin-EDTA plus collagenase 100 mg/mL (Sigma, C9891). Cells were resuspended in PBS/DMEM containing 20% FBS, filtered through a 40 μm mesh to remove debris and cell clumps, stained with DAPI (2 μg/mL), and gated sequentially on cells, singlets, and DAPI-negative live cells before analysis or sorting of DsRed-, mCherry-, or GFP-positive populations based on transgene-associated fluorescence. Fluorescence-activated cell sorting was performed using a BD FACSAria III (BD Biosciences), and analytical flow cytometry was performed using a CytoFLEX S (Beckman Coulter). Flow cytometry data were analyzed using FlowJo v10.10.1.

### Evaluation of endothelial oxidative stress

Endothelial oxidative stress was assessed using the cell-permeant ROS indicator CM-H2DCFDA (Thermo Fisher, C6827). Groups of five embryos were incubated in 1 μg/mL CM-H2DCFDA for 1 h at 28°C in the dark, washed in 1× E3, and dissociated into single-cell suspensions as described in the flow cytometry section. mCherry^+^ endothelial cells were then analyzed by flow cytometry for CM-H2DCFDA fluorescence in the FITC channel after sequential gating on cells, singlets, and DAPI-negative live cells.

### Zebrafish endothelial cell transcriptomic analyses

The transcriptomes of endothelial cells isolated from 24 hpf zebrafish *flk1:mCherry* and *flk1:GFP* embryos were analyzed by RNA-seq. For each experimental group, eight independent pooled single-cell suspensions were generated, each by enzymatic dissociation of two *flk1:mCherry* and two *flk1:GFP* embryos in a single tube. From each mixed preparation, mCherry-positive and GFP-positive endothelial cells were independently isolated by FACS into separate collection tubes, yielding eight sorted mCherry-positive and eight sorted GFP-positive endothelial samples per group, for a total of 16 sorted RNA-seq samples per group, as described in the flow cytometry section. cDNA libraries were constructed from 100–800 collected cells using an in-house adaptation of the Smart-seq2 protocol^86^ in combination with the Nextera XT Library Preparation Kit (Illumina, Inc.). Library size and concentration were evaluated using the TapeStation 2200 system and a Qubit fluorometer before sequencing. Samples were multiplexed, and paired-end 50-bp reads were generated on an Illumina NextSeq 2000 sequencing system. Samples were demultiplexed to generate FASTQ files for each sample. Adapter sequences were removed using Skewer v.0.2.2, and FASTQ files were assessed for quality control using FastQC v.0.11.5. Reads were aligned to the Ensembl zebrafish reference genome GRCz11^87^ using HISAT2 v.2.1.0^88^, with alignment rates greater than 80% for all samples. Gene-level counts were quantified using HTSeq-count v.0.12.4^89^. One sample, grik1b_10, was excluded from downstream analysis after unsupervised principal component analysis identified it as a clear outlier relative to all other samples. Differential gene expression analysis was performed on the remaining samples using DESeq2 v.1.28.1^90^ with genes considered significant at Benjamini–Hochberg-adjusted P < 0.05. Gene ontology and Kyoto Encyclopedia of Genes and Genomes pathway enrichment analyses were performed using GSEA^91^ and clusterProfiler v.4.12.6^92,93^, including gseGO and gseKEGG.

### Mouse and human single-cell transcriptomic analysis

Publicly available mouse single-cell transcriptomic data from the prenatal developmental atlas^43^ were analyzed and visualized using cowplot (v 1.2.0)^94^ and ggplot2 (v 4.0.2)^95^. We selected the annotated cell types Mesodermal_progenitors_Tbx6, Hematoendothelial_progenitors, Arterial_endothelial_cells, Hematopoietic_stem_cells_Cd34, Primitive_erythroid_cells, Border_associated_macrophages, and Glutamatergic_neurons, and restricted the visualization to stages E8.5, E9.5, E10.5, and E11.5. For each selected cell type, dot plots were generated with ggplot2 to show normalized *Grik1* and *Grik5* expression and the percentage of cells expressing each gene. Cell type labels were simplified for visualization.

Publicly available human fetal single-cell RNA-seq data^44^ were analyzed in R using Seurat (v 5.3.0)^96^, tidyverse, readxl, cowplot, and patchwork (v1.3.2)^97^. The published Seurat object (seurat_object.Rdata) was loaded and updated to Seurat v5. For tissue-level visualization, the MAGIC_RNA assay was used to examine *GRIK1* and *GRIK5* expression across AGM, Cord, Head, Heart, Liver, Placenta, VArtery, and YolkSac. Expression values were extracted using FetchData, reshaped for plotting, and visualized as violin plots with overlaid jittered single-cell values. For higher-resolution analysis of hematopoietic emergence, AGM cells were subset from the full object. AGM cluster identities were assigned using marker-defined lineage scoring based on RNA-assay expression values. Marker sets derived from gene scorecards provided in the authors’ shared code were organized into Erythroid, Endothelium, HSPC program, Stroma, Epithelium, Myeloid, General Hematopoietic, and Cycling states, and per-cell scores were calculated as the mean expression of genes within each marker set. Cluster-level mean scores were then used to conservatively assign clusters to broad biological identities, including endothelial-to-hematopoietic transition-like cells (EHT-like), HSPC-like, Erythroid, Myeloid, Stroma, Epithelium, Cycling, and Unknown. After annotation, the assay was switched back to MAGIC_RNA, and *GRIK1* and *GRIK5* expression across AGM cell states was visualized using violin plots.

### Microscopy and time-lapse imaging

Embryos and larvae were mounted for imaging in low-melting-point agarose (Lonza, 50081) containing 0.16 mg/mL tricaine in 35-mm glass-bottom dishes (MatTek) or on microscope slides, and overlaid with E3 medium containing 0.16 mg/mL tricaine. Imaging was performed using an A1R confocal microscope (Nikon), a Nikon 90i upright epifluorescence microscope, a Dragonfly spinning-disk confocal microscope (Andor), or an ImageXpress HCS.ai High-Content Screening System (Molecular Devices). Image brightness and contrast were adjusted to improve visualization.

For time-lapse imaging, embryos were imaged continuously on the Dragonfly spinning-disk confocal microscope using a 20×/0.75 dry objective. Z-stacks were acquired every 5 min for 10 h, with 11 z-planes collected at 5-µm intervals over a 50-µm range. Time-lapse datasets were acquired with two fluorescence channels and processed in Imaris version 9.1. For final visualization, raw 3D time-series datasets were corrected for translational drift using manually placed reference-frame keyframes, displayed as maximum-intensity projections, adjusted for brightness and contrast, and exported as movies with timestamp and scale-bar overlays.

Fluorescence images were quantified from maximum-intensity projections of confocal or epifluorescence z-stacks using Fiji^98^. Cell numbers were determined manually or using semi-automated workflows based on fluorescence thresholding and the Find Maxima function, with analysis parameters applied consistently across directly compared groups.

### EdU staining

At 72 hpf, *runx1:eGFP* embryos were anesthetized in 0.16 mg/mL tricaine and microinjected intravascularly into the duct of Cuvier with 1 nL of 500 μM EdU. After injection, embryos were maintained at 28°C for 1 h, then euthanized and fixed in 4% paraformaldehyde for 1 h, followed by permeabilization in 0.3% Triton X-100 for 20 min at room temperature. EdU incorporation was detected using the Click-iT reaction with Alexa Fluor 647 (Thermo Fisher, C10340). Embryos were then washed in PBS containing 0.3% Triton X-100, blocked for 1 h in 10 mg/mL BSA, 2% FBS, 1% DMSO, and 0.3% Triton X-100, and incubated with anti-GFP 1:200 (Invitrogen, A2311) for 1 h at room temperature. Samples were subsequently washed five times in PBS containing 0.5% Triton X-100, and imaging was performed as described in the microscopy section.

### TUNEL assays

Zebrafish *runx1:eGFP* embryos at 3 dpf were euthanized, fixed, and permeabilized in PBS containing 0.3% Triton X-100 and 1 μg/mL proteinase K for 1 h at room temperature, and TUNEL labeling was performed using the ApopTag Red In Situ Apoptosis Detection Kit (Millipore Sigma, S7165) as previously described^68^. Briefly, embryos were incubated in equilibration buffer for 1 h at room temperature, followed by incubation in TdT reaction mix overnight at 37°C. Larvae were then washed several times in PBST, blocked in 10 mg/mL BSA, 2% FBS, 1% DMSO, and 0.1% Triton X-100, and incubated with anti-digoxigenin-rhodamine (1:200; Roche, 11207750910), and imaging was performed as described in the microscopy section.

### Whole-mount hybridization chain reaction and in situ hybridization assays

Whole-mount hybridization chain reaction was performed on zebrafish embryos as previously described and according to the manufacturer’s protocol^99^ (Molecular Instruments). Embryos were fixed in 4% paraformaldehyde, dehydrated in methanol, and stored at -20°C until processing. Prior to hybridization, embryos were rehydrated, permeabilized, and equilibrated in hybridization buffer. Samples were incubated overnight with the indicated probe sets, washed to remove excess probe, and then incubated overnight with fluorophore-conjugated hairpins that had been snap-cooled before use. After amplification, embryos were washed, mounted, and imaged by confocal microscopy as indicated in the microscopy imaging section.

For whole-mount in situ hybridization (WISH), full-length *runx1* cDNA was cloned into pCRII-TOPO (Invitrogen). Linearized plasmid templates were used to synthesize DIG-labeled antisense RNA probes by *in vitro* transcription with the DIG RNA Labeling Kit (Roche), following the manufacturer’s instructions. The *runx1* probe and WISH protocol have been described previously ^100^, and the probe-cloning primer sequences are listed in Supplementary Table 1.

### Zebrabow color labeling and clonal analysis

At 24 hours post-fertilization (hpf), embryos were transferred to 6-well plates at a density of 25–35 embryos per well and exposed to 15 μM 4-hydroxytamoxifen (4-OHT) for 3–5 h at 28.5°C in the dark. Embryos with weak transgene fluorescence were excluded, and the 25–35 brightest embryos from each condition were retained to minimize variation associated with Zebrabow transgene integration number.

To collect kidney marrows, adult zebrafish (2–9 months old) were anesthetized with 0.02% tricaine and dissected under a Leica MZ75 light microscope. Kidney marrow tissue was placed in cold 0.9x DPBS with 2% fetal bovine serum (Gemini Bio-Products) and 1 USP unit/mL heparin (Sigma), and then mechanically dissociated by repeated pipetting. Samples were passed through a 40-micron nylon mesh 5-10 minutes prior to analysis. We used 3 nM DRAQ-7 for live/dead stain (Abcam). Flow cytometric analysis was performed on a BD FACSAria II with special order 445 nm laser for CFP detection (BD Biosciences). Gates were drawn using negative controls. Flow cytometric data were initially processed in FlowJo software version 10 to extract live granulocytic color output. These cells were chosen for their short half-life and representative clonal output of HSPCs. Color barcodes were quantified using previously published pipelines^4,33^, adapted to a Python-based interface. Only zebrafish with greater than 75% recombination efficiency were processed. All samples were blinded prior to analysis and compared against clutch-matched siblings.

### Phylogenetic analysis of zebrafish KAR subunits

Full-length protein sequences of zebrafish KAR subunits were aligned with MUSCLE ^101^, and a maximum-likelihood phylogenetic tree was constructed in R using phangorn^102^. GluA1a and GluN1a, which are AMPA- and NMDA-type ionotropic glutamate receptor subunits, respectively, were included as outgroups to root the tree. These proteins belong to the same ionotropic glutamate receptor family as KARs but fall outside the kainate receptor subfamily.

### Statistical Analysis

Statistical analyses were performed in GraphPad Prism versions 8.1.2 or 11.0.0 and R version 4.4.1. Unpaired Student’s or Welch’s t tests, Mann–Whitney tests, one- or two-way ANOVA, and Kruskal–Wallis tests were used as appropriate. All tests were two-tailed unless otherwise indicated, and multiple-comparison procedures are specified in the corresponding figure legends. For live-cell quantifications, measurements were normalized to the matched control group within each experiment to account for differences in image acquisition conditions across microscopes and institutions. Sample sizes were based on prior zebrafish studies, preliminary optimization experiments, and the ability to detect reproducible phenotypic differences across independent biological replicates. Except for sequencing experiments and *in situ* hybridization, experiments were performed independently at least twice. Embryos from multiple clutches were pooled before random allocation to control and perturbation groups, whereas Zebrabow clonal-tracing experiments used embryos randomized between conditions within individual clutches. All plotted samples represent independent biological replicates. Unless otherwise specified in the figure legends, n denotes the number of individual larvae analyzed; for pooled assays, n denotes the number of independent pools.

## Data availability

RNA-seq data have been deposited in the NCBI Gene Expression Omnibus and will be publicly available under accession number GSE342053 upon publication: (https://www.ncbi.nlm.nih.gov/geo/query/acc.cgi?acc=GSE342053)

## Acknowledgements

We thank David Brass (Van Andel Institute, VAI) for helpful advice on manuscript preparation, Olivia Olicari (VAI) for assistance with image acquisition and experimental setup on the High-Content Screening System, Rachael Sheridan and the Van Andel Institute Flow Cytometry Core (RRID:SCR_022685) for advice on flow cytometry experimental design, and Senya Combs and Catherine Choi for technical assistance with knockdown reagents. We also acknowledge the VAI Optical Imaging Core (RRID:SCR_021968) for support with microscopy and image acquisition. This research was supported in part by VAI‘s West Michigan Neurodegenerative Diseases (MiND) Program iPSC and Screening Platform (RRID:SCR_027675). AI-assisted tools were used for language editing. Figures were partially created using BioRender. Parada Kusz, M. (2026). BioRender.com/3esaf4k.

## Author contributions

Study conception and design were led by M.P.-K., with contributions from S.W., A.E.C., S.G., and D.T.H. Experiments and data analysis were performed by M.P.-K., S.W., J.J., J.S., A.I., V.d.N., E.G., S.B., M.E., Z.F., A.E.C., and T.W. The manuscript was written by M.P.-K. and revised with contributions from S.W., T.W., A.E.C., S.G., and D.T.H.

---

**Supplementary Figure 1. KAR subunit organization and validation of CRISPR-mediated gene disruption.**

**a** Domain organization of all kainate receptor subunits encoded in the zebrafish genome. Schematics show each protein alongside its corresponding gene name and depict the amino-terminal domain (ATD), ligand-binding domain (LBD), and transmembrane segments (TM), aligned by residue position for comparison. **b** Maximum-likelihood tree of zebrafish KARs based on MUSCLE alignment of full-length amino acid sequences. AMPA and NMDA receptor subunits GluA1a and GluN1a, which belong to the broader ionotropic glutamate receptor family, were included as outgroups. **c** Experimental workflow for quantifying innate immune cells in *mpeg1:mCherry;mpx:GFP* larvae. KAR function was reduced by CRISPR- or morpholino-mediated (MO) knockdown at the one-cell stage, or by continuous treatment with KAR inhibitors from 5 to 72 hpf, followed by live imaging of macrophages and neutrophils at 72 hpf. **d-f** CRISPR/Cas9-induced mutations in *grik1a* **d**, *grik1b* **e**, and *grik5* **f** in F0 crispants. CRISPR/Cas9 reagents targeting the indicated exons, with the corresponding guide identifiers shown above each plot, were injected into 1-cell-stage embryos. At 72 hours post-fertilization, larvae from CRISPR/Cas9-injected and uninjected groups were collected for genomic DNA isolation, and regions surrounding each guide RNA target site were amplified by PCR. Amplicons from individual fish were analyzed by paired-end next-generation sequencing to quantify editing efficiency at each locus. Sequence reads were analyzed using CRISPResso2, and the percentages of unmodified and modified reads are shown for one representative larva, with the corresponding percentages indicated above the bars. **g**-**i** RT-qPCR analysis of *grik1a* **g**, *grik1b* **h**, and *grik5* **i** mRNA expression in crispants relative to uninjected controls. **g**, **h**: n = 3 biological replicates; **i**: n = 4 biological replicates, each consisting of 8 fish per group from an independent clutch; individual data points are overlaid. Statistics: panel **g**, one-tailed Mann–Whitney test; panel **h**, unpaired Student’s t test; panel **i**, Mann–Whitney test.

**Supplementary Figure 2. KAR deficiency reduces innate immune cell abundance while preserving early myeloid populations.**

**a** Representative whole-larva epifluorescence images of control and *pu.1* morphant larvae at 72 hpf. Macrophages are pseudocolored magenta and neutrophils green. **b**, **c** Relative macrophage and neutrophil counts, respectively, in control and *pu.1* morphant larvae at 72 hpf. **d**, **e** Relative macrophage and neutrophil counts, respectively, in control and *grik1b* morphant larvae at 72 hpf. **f**, **g** Relative macrophage and neutrophil counts, respectively, in control and *grik5* morphant larvae at 72 hpf. **h**, **i** Relative macrophage and neutrophil counts, respectively, in control and *grik1a* CRISPR larvae at 72 hpf. **j** Representative confocal maximum-intensity projections showing whole-mount hybridization chain reaction detection of *pu.1* expression in control, *grik1b*, and *grik5* morphants at 24 hpf. Dashed line indicates the approximate boundary used to distinguish anterior and posterior myeloid cell populations. **k**, **l** Quantification of relative anterior **k** and posterior **l** *pu.1+* cells at 24 hpf. Data are shown as box-and-whisker plots with individual values; center line, median; whiskers, minimum to maximum. Statistics: panels **b** and **c**, one-way ANOVA with Dunnett’s multiple-comparisons test; panels **d**, **g**, **h**, and **i**, Welch’s unpaired t test; panels **e** and **f**, Mann–Whitney test; panels **k** and **l**, Kruskal–Wallis test with Dunn’s multiple-comparisons test.

**Supplementary Figure 3. KAR deficiency impairs primitive erythropoiesis without overt endothelial defects at 24 hpf.**

**a** Representative flow cytometry plots of dissociated *gata1a* larvae at 72 hpf showing the percentage of DsRed^+^ erythrocytes in control, *grik1b*, and *grik5* morphants. Cells were sequentially gated on total cells, singlets, and live cells before assessment of DsRed fluorescence. **b** Representative lateral images of *flk1* embryos at 24 hpf showing endothelial fluorescence and corresponding bright-field views in control, *grik1b*, and *grik5* morphants. **c** Representative flow cytometry plots of single-cell suspensions prepared from *flk1* embryos at 24 hpf showing the fraction of mCherry+ endothelial cells in control, *grik1b*, and *grik5* morphants. **d** Quantification of the percentage of mCherry+ cells from flow cytometry analysis. Each dot in **d** represents one biological replicate composed of five pooled embryos. **e, f** Quantification of heartbeat rate (**e**) and the number of circulating blood cells passing a defined location in the dorsal aorta per 15 s (**f**) at 26 hpf in control, *grik1b*, and *grik5* morphants. Data are shown as box-and-whisker plots with individual values; center line, median; whiskers, min to max. Statistics: panel **d**, one-way ANOVA with Dunnett’s multiple-comparisons test; panels **e** and **f**, Kruskal–Wallis test with Dunn’s multiple-comparisons test.

**Supplementary Figure 4. Evaluation of HSPC proliferation, survival, and clonal-tracing strategy.**

**a** Schematic showing the developmental timing of macrophage-dependent regulation of HSPC emergence in the hemogenic niche and subsequent HSPC expansion/attrition in the caudal hematopoietic tissue. **b** Representative confocal maximum-intensity projections of the caudal hematopoietic tissue at 72 hpf in *flk1:mCherry;runx1:eGFP* larvae showing reduced HSPC abundance in *grik1b* and *grik5* morphants. Endothelial cells are pseudocolored magenta. **c** Representative confocal maximum-intensity projections of the caudal hematopoietic tissue at 72 hpf in *runx1:eGFP* larvae following control, *grik1b*, or *grik5* morpholino injection and TUNEL labeling, showing no increase in TUNEL+ HSPCs following KAR knockdown. **d** Quantification of the percentage of TUNEL+ HSPCs in the caudal hematopoietic tissue at 72 hpf. **e** Representative confocal maximum-intensity projections of the caudal hematopoietic tissue at 72 hpf in *runx1:eGFP* larvae following control, *grik1b*, or *grik5* morpholino injection and EdU labeling, showing reduced EdU incorporation in HSPCs following KAR knockdown. Arrowheads indicate representative EdU+ HSPCs. **f** Quantification of the percentage of EdU+ HSPCs in the caudal hematopoietic tissue at 72 hpf in control, *grik1b*, and *grik5* morphants. **g** Schematic of the Zebrabow-M clonal-tracing strategy. Fish carrying 15–20 integrations of the multicolor Zebrabow cassette were crossed with *drl:CreERT2* animals, and embryos were injected with control, *grik1b*, or *grik5* morpholinos. Treatment with 4-hydroxytamoxifen (4-OHT) at 24 hpf induced stochastic Cre-mediated recombination in drl-derived lateral plate mesoderm lineages, generating heritable fluorescent hues that were quantified in adult kidney marrow to estimate HSC clonal contributions. Statistics: panel **d**, Kruskal–Wallis test with Dunn’s multiple-comparisons test; panel **f**, one-way ANOVA with Holm–Šídák’s multiple-comparisons test. **h** Representative ternary plots showing Zebrabow fluorescence profiles of adult kidney marrow granulocytes from fish injected with control, *grik1b*, or *grik5* morpholinos during embryogenesis. Each point represents one cell positioned according to the relative contributions of RFP, CFP, and YFP fluorescence. Each plot represents one adult fish and illustrates the color-barcode distributions underlying the clonal quantification.

**Supplementary Figure 5. *gata1a* co-knockdown restores whole-larva macrophage and neutrophil abundance and HSPC numbers in the caudal hematopoietic tissue of KAR-deficient larvae.**

**a**, **b** Relative macrophage counts in whole larvae at 72 hpf following *grik1b* or *grik5* morpholino knockdown, *gata1a* morpholino knockdown, or combined KAR/*gata1a* knockdown, compared with controls. **c**, **d** Relative neutrophil counts in whole larvae at 72 hpf following *grik1b* or *grik5* morpholino knockdown, *gata1a* morpholino knockdown, or combined KAR/*gata1a* knockdown, compared with controls. **e**, **f** Relative HSPC counts in the caudal hematopoietic tissue at 72 hpf following *grik1b* or *grik5* morpholino knockdown, alone or in combination with *gata1a* morpholino, compared with controls. Data are shown as box-and-whisker plots with individual values; center line, median; whiskers, min to max. Statistics: panels **a**, **b**, **e**, and **f**, one-way ANOVA with Šidák’s multiple-comparisons test; panel **c**, one-way ANOVA with Dunnett’s multiple-comparisons test; panel **d**, Kruskal–Wallis test with Dunn’s multiple-comparisons test.

**Supplementary Figure 6. Endothelial transcriptomic profiling identifies stress-associated and developmental pathway changes following KAR knockdown.**

**a** Experimental design for endothelial RNA-seq. *flk1:mCherry* and *flk1:GFP* embryos injected with control, *grik1b*, or *grik5* morpholino (MO) were collected at 24 hpf, dissociated, and endothelial cells were isolated by FACS for RNA-seq and differential-expression analysis. **b, c** GSEA summary plots showing selected GO biological processes and KEGG signaling pathways in *grik1b* and *grik5* morphants, each relative to controls. Dot position indicates normalized enrichment score (NES), dot size indicates gene-set size, and color indicates false discovery rate (FDR). FoxO signaling trended toward positive enrichment in *grik1b* morphants **d** and was significantly enriched in *grik5* morphants **e**; cell cycle **f** and blood circulation **g** were negatively enriched, whereas blood vessel development **h** and Notch signaling **i** were positively enriched in *grik5* morphants. **j** Representative flow cytometry histogram overlays showing CM-H2DCFDA fluorescence in mCherry+ endothelial cells from dissociated *flk1:mCherry* embryos at 24 hpf in control, *grik1b*, and *grik5* morphants. **k** Representative confocal maximum-intensity projections of *fli1a:GFP;Tp1:mCherry* embryos at 30 hpf showing Notch-responsive endothelial cells in control, *grik1b*, *grik5*, *grik1b* + *gata1a*, and *grik5* + *gata1a* morphants. Tp1+ cells are pseudocolored magenta. Scale bar, 20 μm. **l** Representative *runx1* whole-mount in situ hybridization at 30 hpf in control, *grik1b*, and *grik5* morphants with or without *gata2b:KalTA4;UAS:NICD*-mediated constitutive Notch activation. NES, normalized enrichment score; FDR, false discovery rate.

**Supplementary Figure 7. KAR subunit expression is conserved across neural, endothelial, and hematopoietic populations during vertebrate development.**

**a** UMAP of the published zebrafish whole-embryo single-cell RNA-seq atlas showing annotated cell populations from *Sur et al., 2023*^42^. **b**, **c** Feature plots showing *grik1b* **b** and *grik5* c expression across the zebrafish atlas. Expression is most prominent in neural populations, with additional signal in vascular, immune, and erythroid clusters, particularly for *grik5*. **d** Normalized RNA-seq expression of *grik1b*, *grik5*, and *flk1* in zebrafish endothelial cells isolated at 24 hpf. **e** Dot plots showing *Grik1* and *Grik5* expression across mesodermal, hematoendothelial, endothelial, hematopoietic, and neuronal populations in mouse embryos from E8.5 to E11.5, based on reanalysis of the published scRNA-seq dataset from *Qiu et al., 2024*^43^. Dot size indicates the fraction of expressing cells, and color indicates average expression level.

**Supplementary Movies 1–3**. Live imaging of HSPC emergence in control and KAR-deficient embryos.

Time-lapse spinning-disk confocal imaging of *flk1:mCherry;runx1:eGFP* embryos beginning at 30 hpf and continuing for 10 h. Movies show merged maximum-intensity projections of z-stacks through the dorsal aorta region in control embryos, *grik1b* morphants, and *grik5* morphants, illustrating reduced HSPC emergence in KAR-deficient embryos. The mCherry signal marking endothelial cells is pseudocolored magenta.

