## Supplementary Figures for "Kainate receptors coordinate primitive hematopoiesis and hemogenic niche organization to promote hematopoietic stem cell development"

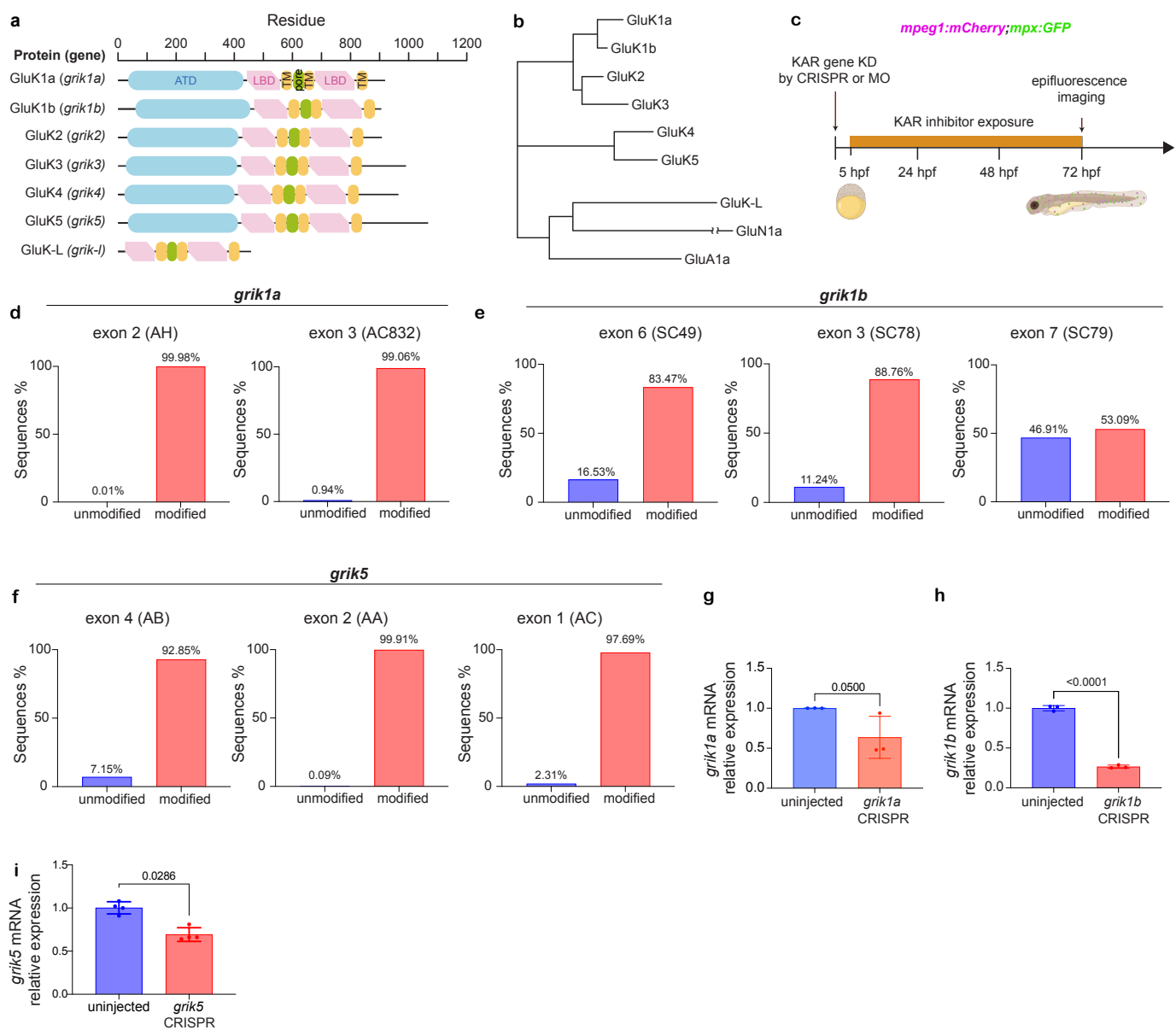

### Supplementary Figure 1. KAR subunit organization and validation of CRISPR-mediated gene disruption.

**a** Domain organization of all kainate receptor subunits encoded in the zebrafish genome. Schematics show each protein alongside its corresponding gene name and depict the amino-terminal domain (ATD), ligand-binding domain (LBD), and transmembrane segments (TM), aligned by residue position for comparison. **b** Maximum-likelihood tree of zebrafish KARs based on MUSCLE alignment of full-length amino acid sequences. AMPA and NMDA receptor subunits GluA1a and GluN1a, which belong to the broader ionotropic glutamate receptor family, were included as outgroups. **c** Experimental workflow for quantifying innate immune cells in *mpeg1:mCherry;mpx:GFP* larvae. KAR function was reduced by CRISPR- or morpholino-mediated (MO) knockdown at the one-cell stage, or by continuous treatment with KAR inhibitors from 5 to 72 hpf, followed by live imaging of macrophages and neutrophils at 72 hpf. **d-f** CRISPR/Cas9-induced mutations in *grik1a* **d**, *grik1b* **e**, and *grik5* **f** in F0 crispants. CRISPR/Cas9 reagents targeting the indicated exons, with the corresponding guide identifiers shown above each plot, were injected into 1-cell-stage embryos. At 72 hours post-fertilization, larvae from CRISPR/Cas9-injected and uninjected groups were collected for genomic DNA isolation, and regions surrounding each guide RNA target site were amplified by PCR. Amplicons from individual fish were analyzed by paired-end next-generation sequencing to quantify editing efficiency at each locus. Sequence reads were analyzed using CRISPResso2, and the percentages of unmodified and modified reads are shown for one representative larva, with the corresponding percentages indicated above the bars. **g-i** RT-qPCR analysis of *grik1a* **g**, *grik1b* **h**, and *grik5* **i** mRNA expression in crispants relative to uninjected controls. **g,h**: n = 3 biological replicates; **i**: n = 4 biological replicates, each consisting of 8 fish per group from an independent clutch; individual data points are overlaid. Statistics: panel **g**, one-tailed Mann–Whitney test; panel **h**, unpaired Student's t test; panel **i**, Mann–Whitney test.

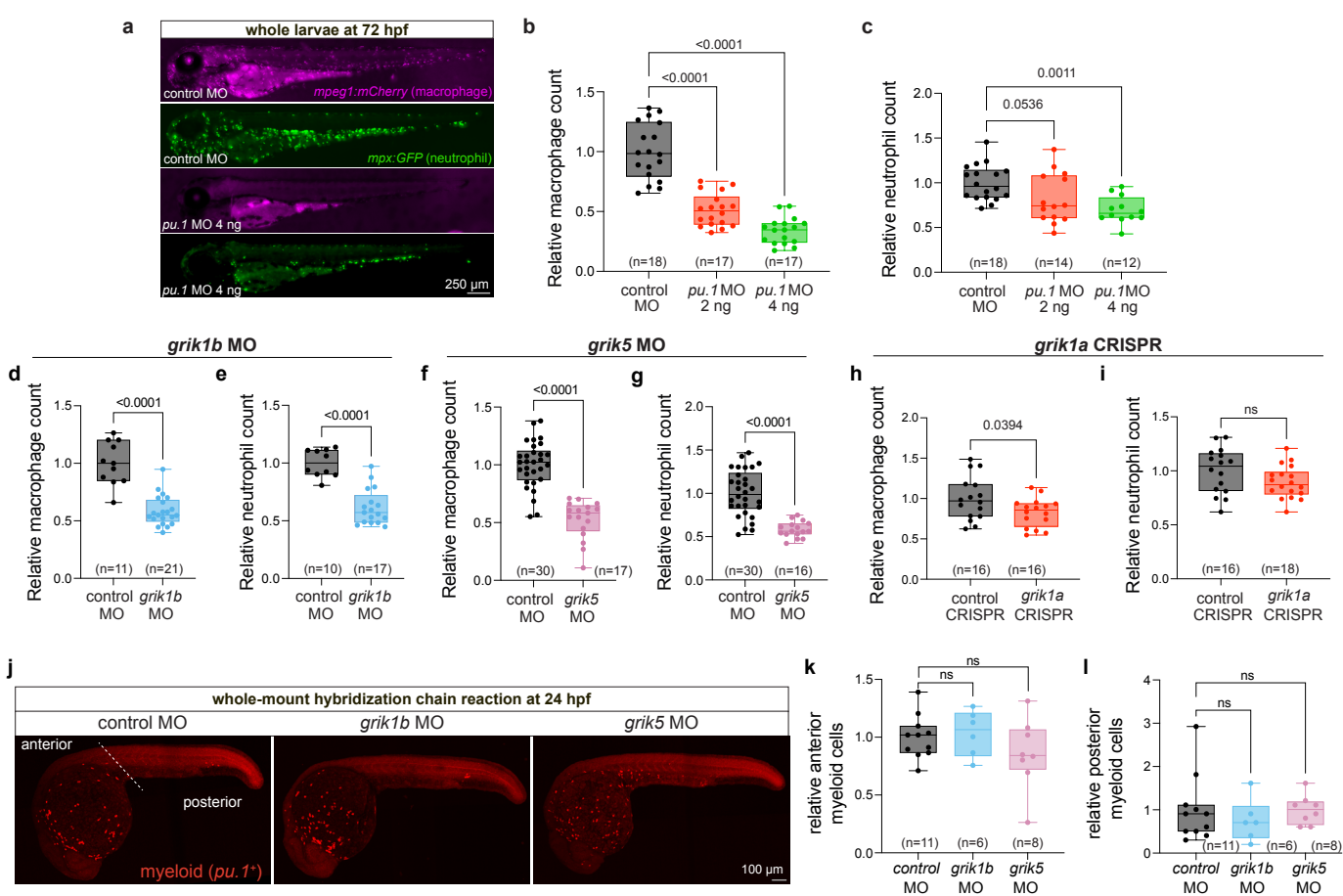

**Supplementary Figure 2. KAR deficiency reduces innate immune cell abundance while preserving early myeloid populations.**

**a** Representative whole-larva epifluorescence images of control and *pu.1* morphant larvae at 72 hpf. Macrophages are pseudocolored magenta. **b, c** Relative macrophage and neutrophil counts, respectively, in control and *pu.1* morphant larvae at 72 hpf. **d, e** Relative macrophage and neutrophil counts, respectively, in control and *grik1b* morphant larvae at 72 hpf. **f, g** Relative macrophage and neutrophil counts, respectively, in control and *grik5* morphant larvae at 72 hpf. **h, i** Relative macrophage and neutrophil counts, respectively, in control and *grik1a* CRISPR larvae at 72 hpf. **j** Representative confocal maximum-intensity projections showing whole-mount hybridization chain reaction detection of *pu.1* expression in control, *grik1b*, and *grik5* morphants at 24 hpf. Dashed line indicates the approximate boundary used to distinguish anterior and posterior myeloid cell populations. **k, l** Quantification of relative anterior **k** and posterior **l** *pu.1*<sup>+</sup> cells at 24 hpf. Data are shown as box-and-whisker plots with individual values; center line, median; whiskers, minimum to maximum. Statistics: panels **b** and **c**, one-way ANOVA with Dunnett's multiple-comparisons test; panels **d, g, h**, and **i**, Welch's unpaired t test; panels **e** and **f**, Mann–Whitney test; panels **k** and **l**, Kruskal–Wallis test with Dunn's multiple-comparisons test.

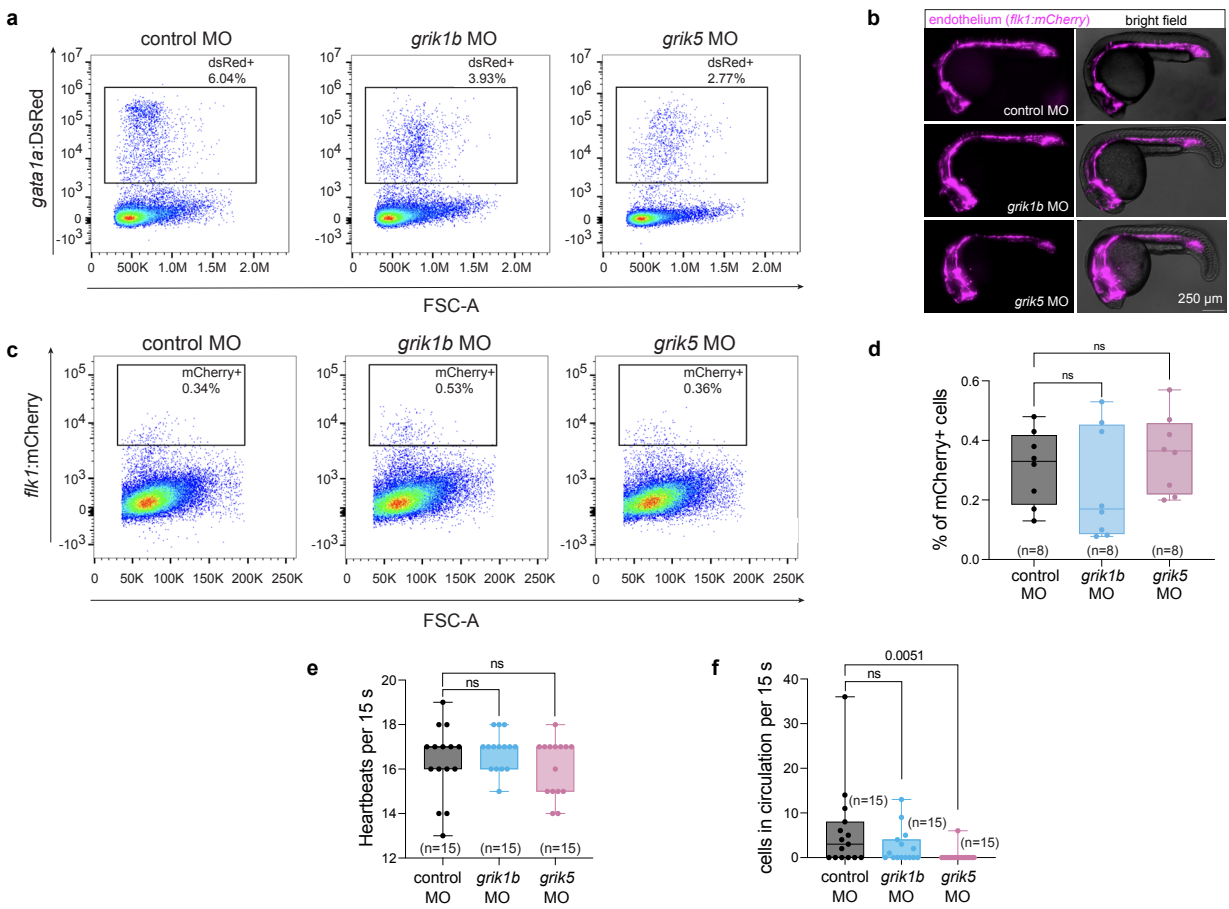

**Supplementary Figure 3. KAR deficiency impairs primitive erythropoiesis without overt endothelial defects at 24 hpf.**

**a** Representative flow cytometry plots of dissociated *gata1a* larvae at 72 hpf showing the percentage of DsRed<sup>+</sup> erythrocytes in control, *grik1b*, and *grik5* morphants. Cells were sequentially gated on total cells, singlets, and live cells before assessment of DsRed fluorescence. **b** Representative lateral images of *flk1* embryos at 24 hpf showing endothelial fluorescence and corresponding bright-field views in control, *grik1b*, and *grik5* morphants. **c** Representative flow cytometry plots of single-cell suspensions prepared from *flk1* embryos at 24 hpf showing the fraction of mCherry<sup>+</sup> endothelial cells in control, *grik1b*, and *grik5* morphants. **d** Quantification of the percentage of mCherry<sup>+</sup> cells from flow cytometry analysis. Each dot in **d** represents one biological replicate composed of five pooled embryos. **e**, **f** Quantification of heartbeat rate (**e**) and the number of circulating blood cells passing a defined location in the dorsal aorta per 15 s (**f**) at 26 hpf in control, *grik1b*, and *grik5* morphants. Data are shown as box-and-whisker plots with individual values; center line, median; whiskers, min to max. Statistics: panel **d**, one-way ANOVA with Dunnett's multiple-comparisons test; panels **e** and **f**, Kruskal–Wallis test with Dunn's multiple-comparisons test.

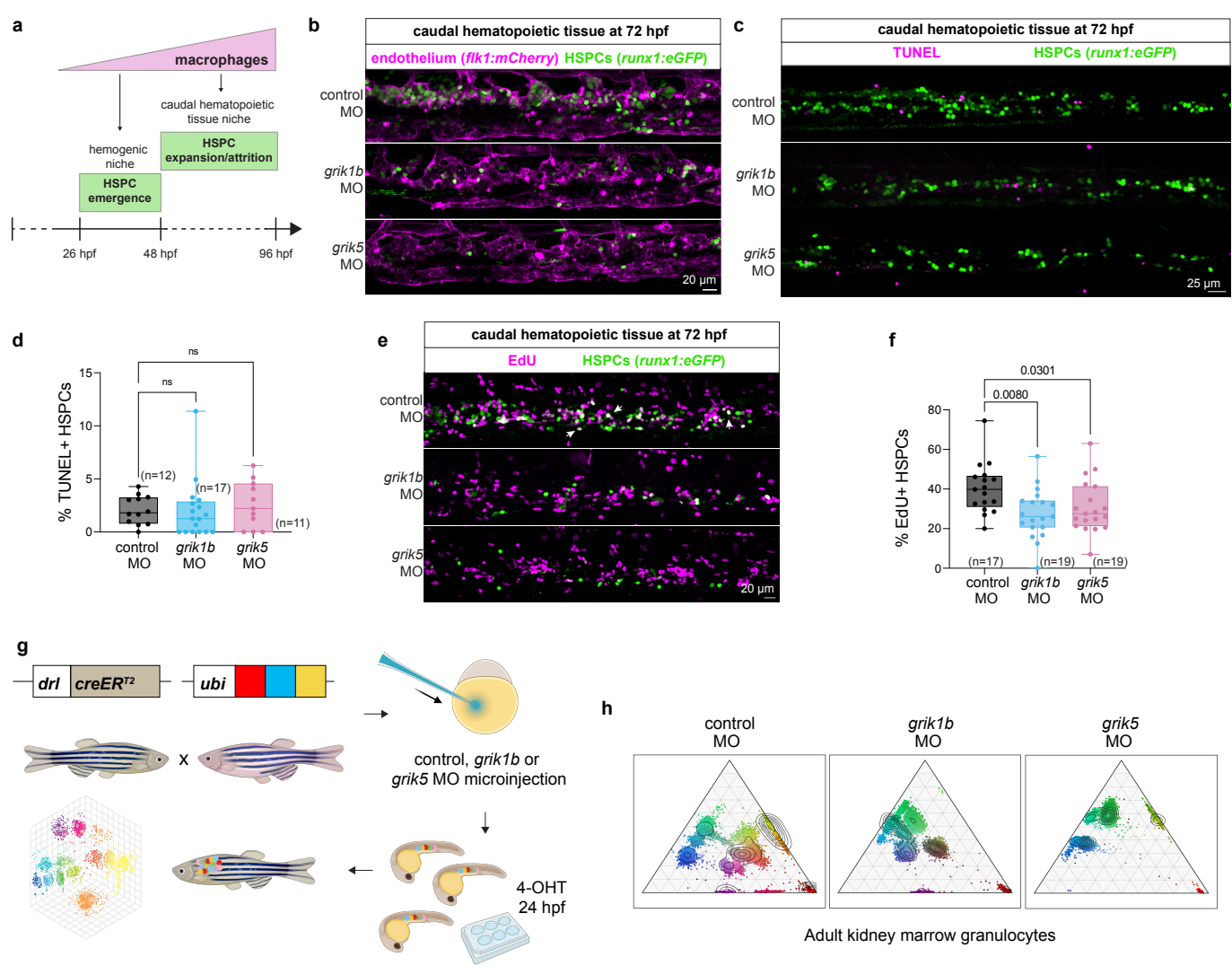

**Supplementary Figure 4. Evaluation of HSPC proliferation, survival, and clonal-tracing strategy.**

**a** Schematic showing the developmental timing of macrophage-dependent regulation of HSPC emergence in the hemogenic niche and subsequent HSPC expansion/attrition in the caudal hematopoietic tissue. **b** Representative confocal maximum-intensity projections of the caudal hematopoietic tissue at 72 hpf in *flk1:mCherry*; *runx1:eGFP* larvae showing reduced HSPC abundance in *grik1b* and *grik5* morphants. Endothelial cells are pseudocolored magenta. **c** Representative confocal maximum-intensity projections of the caudal hematopoietic tissue at 72 hpf in *runx1:eGFP* larvae following control, *grik1b*, or *grik5* morpholino injection and TUNEL labeling, showing no increase in TUNEL+ HSPCs following KAR knockdown. **d** Quantification of the percentage of TUNEL+ HSPCs in the caudal hematopoietic tissue at 72 hpf. **e** Representative confocal maximum-intensity projections of the caudal hematopoietic tissue at 72 hpf in *runx1:eGFP* larvae following control, *grik1b*, or *grik5* morpholino injection and EdU labeling, showing reduced EdU incorporation in HSPCs following KAR knockdown. Arrowheads indicate representative EdU+ HSPCs. **f** Quantification of the percentage of EdU+ HSPCs in the caudal hematopoietic tissue at 72 hpf in control, *grik1b*, and *grik5* morphants. **g** Schematic of the Zebrafish-M clonal-tracing strategy. Fish carrying 15–20 integrations of the multicolor Zebrafish cassette were crossed with *drl:CreERT2* animals, and embryos were injected with control, *grik1b*, or *grik5* morpholinos. Treatment with 4-hydroxytamoxifen (4-OHT) at 24 hpf induced stochastic Cre-mediated recombination in *drl*-derived lateral plate mesoderm lineages, generating heritable fluorescent hues that were quantified in adult kidney marrow to estimate HSC clonal contributions. **h** Representative ternary plots showing Zebrafish fluorescence profiles of adult kidney marrow granulocytes from fish injected with control, *grik1b*, or *grik5* morpholinos during embryogenesis. Each point represents one cell positioned according to the relative contributions of RFP, CFP, and YFP fluorescence. Each plot represents one adult fish and illustrates the color-barcode distributions underlying the clonal quantification.

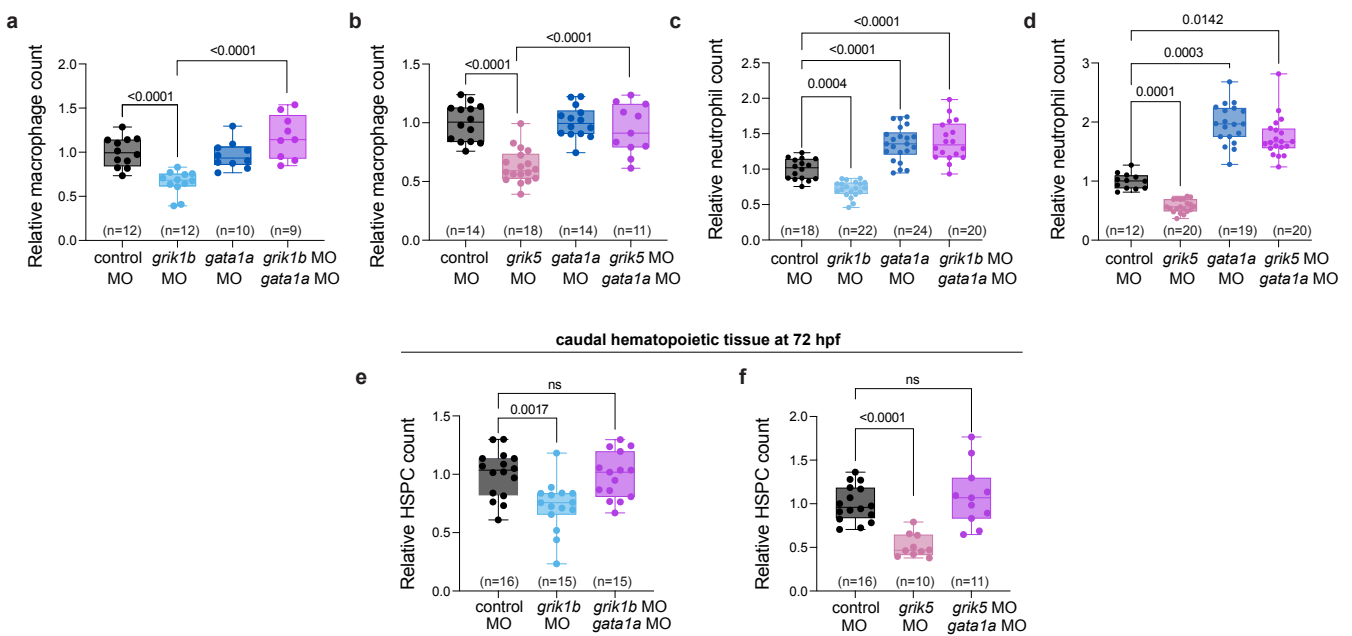

**Supplementary Figure 5. *gata1a* co-knockdown restores whole-larva macrophage and neutrophil abundance and HSPC numbers in the caudal hematopoietic tissue of KAR-deficient larvae.**

**a, b** Relative macrophage counts in whole larvae at 72 hpf following *grik1b* or *grik5* morpholino knockdown, *gata1a* morpholino knockdown, or combined KAR/*gata1a* knockdown, compared with controls. **c, d** Relative neutrophil counts in whole larvae at 72 hpf following *grik1b* or *grik5* morpholino knockdown, *gata1a* morpholino knockdown, or combined KAR/*gata1a* knockdown, compared with controls. **e, f** Relative HSPC counts in the caudal hematopoietic tissue at 72 hpf following *grik1b* or *grik5* morpholino knockdown, alone or in combination with *gata1a* morpholino, compared with controls. Data are shown as box-and-whisker plots with individual values; center line, median; whiskers, min to max. Statistics: panels **a, b, e, and f**, one-way ANOVA with Šídák's multiple-comparisons test; panel **c**, one-way ANOVA with Dunnett's multiple-comparisons test; panel **d**, Kruskal-Wallis test with Dunn's multiple-comparisons test.

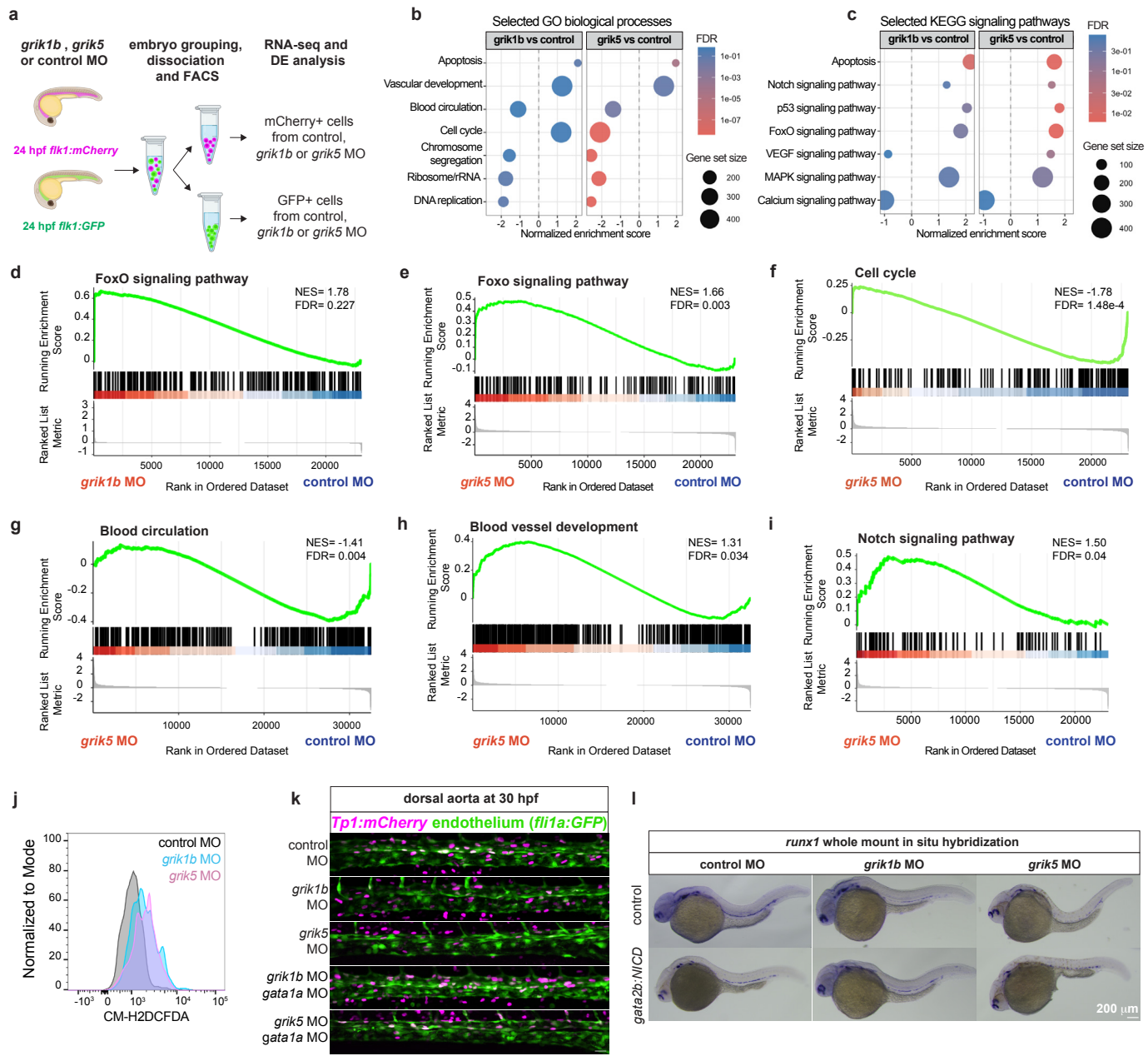

**Supplementary Figure 6. Endothelial transcriptomic profiling identifies stress-associated and developmental pathway changes following KAR knockdown.**

**a** Experimental design for endothelial RNA-seq. *flk1:mCherry* and *flk1:GFP* embryos injected with control, *grik1b*, or *grik5* morpholino (MO) were collected at 24 hpf, dissociated, and endothelial cells were isolated by FACS for RNA-seq and differential-expression analysis. **b**, **c** GSEA summary plots showing selected GO biological processes and KEGG signaling pathways in *grik1b* and *grik5* morphants, each relative to controls. Dot position indicates normalized enrichment score (NES), dot size indicates gene-set size, and color indicates false discovery rate (FDR). FoxO signaling trended toward positive enrichment in *grik1b* morphants **d** and was significantly enriched in *grik5* morphants **e**; cell cycle **f** and blood circulation **g** were negatively enriched, whereas blood vessel development **h** and Notch signaling **i** were positively enriched in *grik5* morphants. **j** Representative flow cytometry histogram overlays showing CM-H2DCFDA fluorescence in mCherry<sup>+</sup> endothelial cells from dissociated *flk1:mCherry* embryos at 24 hpf in control, *grik1b*, and *grik5* morphants. **k** Representative confocal maximum-intensity projections of *flk1a:GFP*, *Tp1:mCherry* embryos at 30 hpf showing Notch-responsive endothelial cells in control, *grik1b*, *grik5*, *grik1b* + *gata1a*, and *grik5* + *gata1a* morphants. Tp1<sup>+</sup> cells are pseudocolored magenta. Scale bar, 20 μm. **l** Representative *runx1* whole-mount in situ hybridization at 30 hpf in control, *grik1b*, and *grik5* morphants with or without *gata2b:KaItA4;UAS:NICD*-mediated constitutive Notch activation. NES, normalized enrichment score; FDR, false discovery rate.

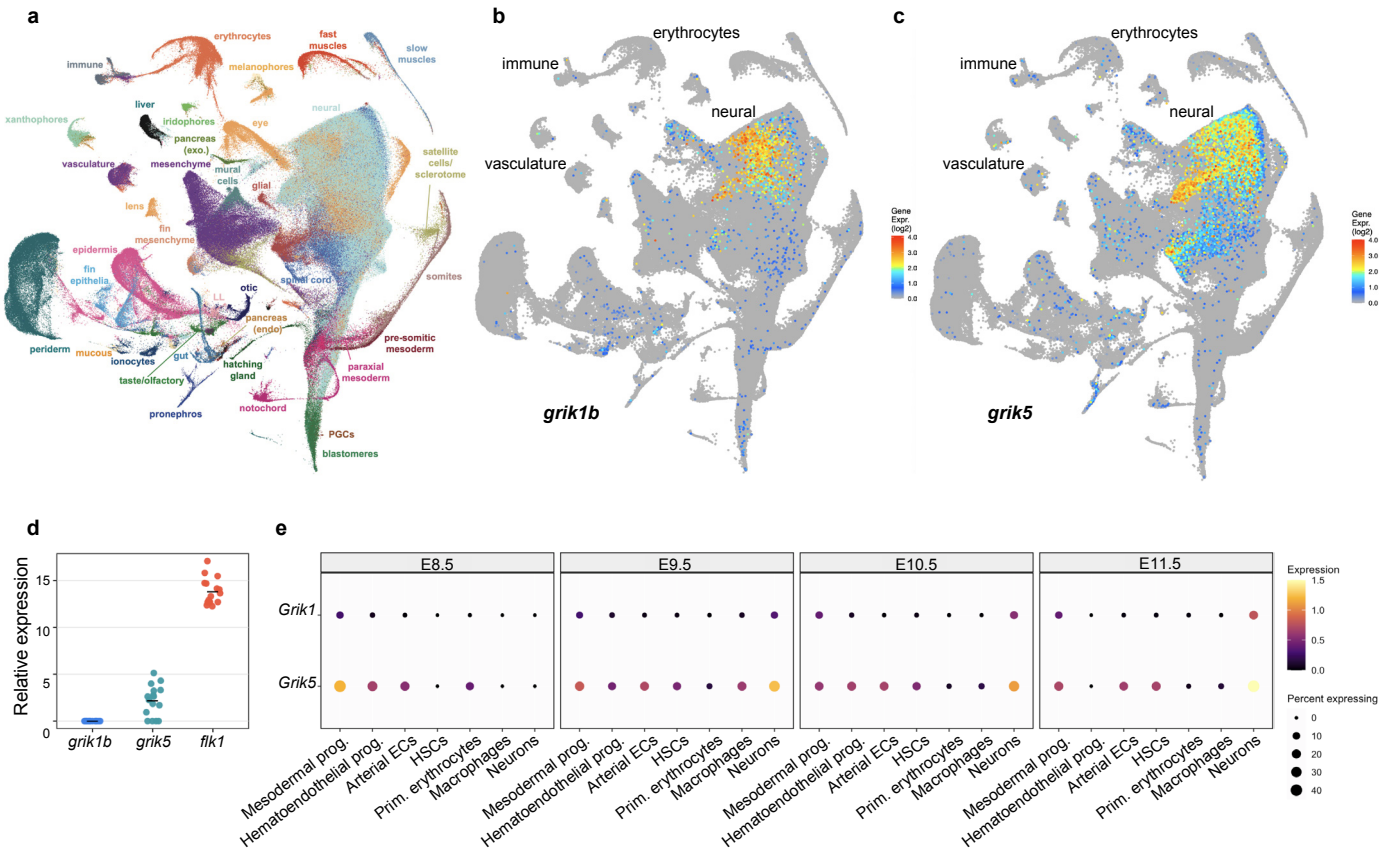

**Supplementary Figure 7. KAR subunit expression is conserved across neural, endothelial, and hematopoietic populations during vertebrate development.**

**a** UMAP of the published zebrafish whole-embryo single-cell RNA-seq atlas showing annotated cell populations from *Sur et al., 2023*<sup>42</sup>. **b, c** Feature plots showing *grik1b* **b** and *grik5* **c** expression across the zebrafish atlas. Expression is most prominent in neural populations, with additional signal in vascular, immune, and erythroid clusters, particularly for *grik5*. **d** Normalized RNA-seq expression of *grik1b*, *grik5*, and *flk1* in zebrafish endothelial cells isolated at 24 hpf. **e** Dot plots showing *Grik1* and *Grik5* expression across mesodermal, hematoendothelial, endothelial, hematopoietic, and neuronal populations in mouse embryos from E8.5 to E11.5, based on reanalysis of the published scRNA-seq dataset from *Qiu et al., 2024*<sup>43</sup>. Dot size indicates the fraction of expressing cells, and color indicates average expression level.
